# Brain proteinopathies associated physiological changes leading to cognitive resilience in Alzheimer’s disease

**DOI:** 10.64898/2026.11.09.765496

**Authors:** Purba Mandal, Priyanka Baloni

## Abstract

Alzheimer’s disease (AD) pathology has been well characterized, however the molecular mechanisms underlying resilience to the disease remain understudied. With the growing evidence of individuals who are resilient to AD and generation of cross-cohort multiomics data, it has become feasible to understand these mechanisms. In this study, we integrated longitudinal cognitive assessments, regional neuropathology, and matched prefrontal proteomic and metabolomic profiles across donors from two independent cohorts. We identified that asymptomatic AD (AsymAD) brains exhibited amyloid burdens comparable to AD but had fewer compact plaques, lower regional tau burden, and more restricted TDP-43 pathology. Across matched stages of amyloid, tau, and TDP-43 pathology, AsymAD was consistently distinguished from AD by preservation of mitochondrial translation and bioenergetics, synaptic programs, glutamatergic metabolites, and polyunsaturated membrane lipids. Longitudinal cognitive assessments further separated resilient from vulnerable AsymAD individuals despite comparable amyloid and TDP-43 burden, with vulnerability associated with greater neurofibrillary tangle accumulation. Both AsymAD groups retained a prefrontal proteome close to that of cognitively normal individuals, apart from amyloid-associated proteins, whereas mitochondrial, synaptic, and immune changes were largely confined to AD.

**Highlights:**

- AsymAD carries AD-level amyloid with lesser compact plaques, tau, and TDP-43
- The mitochondrial, lipid, synaptic functions separate AsymAD from AD at every pathology stage
- Resilient and vulnerable AsymAD differ in tangles, not amyloid or TDP-43
- AsymAD keeps a control-like prefrontal proteome

## Introduction

Alzheimer’s disease (AD) is pathologically characterized by two hallmark lesions, amyloid-β plaques and tau neurofibrillary tangles^1^, yet it is uncommon for the aging brain to present with these lesions in isolation. Most individuals with AD neuropathologic change also accumulate additional proteinopathies, most commonly TAR DNA-binding protein 43 (TDP-43) in the form of limbic-predominant age-related TDP-43 encephalopathy neuropathologic change (LATE-NC), together with α-synuclein, vascular, and other age-related pathologies^2,3^. These co-pathologies accelerate the onset of dementia and are responsible for a large share of patient’s cognitive decline^4,5^. This heterogeneity in co-pathologies influences the therapeutic consequences. Anti-amyloid antibodies such as lecanemab and donanemab slow decline in early symptomatic AD, but their benefit is modest and accompanied by amyloid-related imaging abnormalities^6–8^. Amyloid therefore explains only part of who develops dementia, and the other determinants of clinical expression remain poorly understood.

A subset of older adults in human cohort studies offers potential opportunities to isolate these determinants. These individuals, described as asymptomatic AD (AsymAD), accumulate substantial amyloid and tau pathology yet remain cognitively intact during life. They represent a considerable fraction of autopsied older adults in community-based cohorts, including the Nun Study, the Honolulu-Asia Aging Study, and the Rush cohorts^9–11^, and they represent cognitive resilience, the capacity to maintain cognition despite pathology, which is distinct from resistance, the avoidance of pathology altogether^12^. Postmortem studies have linked resilience to preserved synaptic density, limited glial activation, and neuronal hypertrophy^10,13,14^. Large-scale proteomics has since identified AsymAD as a molecularly distinct state^15–17^ and implicated synaptic proteins such as neuritin (NRN1) in resilience to amyloid-β^18,19^. In our previous work, AsymAD brains maintained higher mitochondrial bioenergetic capacity than AD brains^20^.

These studies establish molecular correlates of resilience yet leave critical gaps in our understanding. The first is whether resilient individuals tolerate the same pathology that causes dementia in others or is their pathology distinct in its type, composition, or accompanying condition. Most studies, including our own, match AsymAD and AD on amyloid and tau stages, and these staging metrics condense plaque subtypes, spatial distribution, and co-pathology into a single score. This oversimplification masks individual variations in pathology. TDP-43 is a strong candidate for a hidden variable because it is present in a large fraction of AD brains, drives cognitive impairment independently of amyloid and tau and cannot currently be detected in living patients^2,21,22^. The second question is whether the molecular programs associated with resilience are a general response to pathology or are tailored to each proteinopathy and its stage. A protective program should persist when AsymAD and AD are compared within matched strata of amyloid, tau, and TDP-43, which separates protective biology from differences in pathological severity. The third question is what distinguishes AsymAD individuals who remain cognitively intact from those who show decline. Some individuals with AsymAD develop mild cognitive impairment (MCI) before death, and comparing them with those who remain unimpaired could identify the changes that indicate the loss of resilience.

Molecular profiling of human brain is possible only after death, so each donor is measured at a single time point. In this study we used longitudinal cognitive scores collected up to 20 years before death, together with clinical diagnoses and autopsy neuropathology, to group donors by cognitive and pathological trajectory. We analyzed two complementary human cohorts to study the molecular signatures associated with cognitive changes in the individuals. The Religious Orders Study and Rush Memory and Aging Project (ROSMAP)^23^ enrolled older adults without known dementia who underwent annual cognitive assessment and their brains were subjected to neuropathological and multi-omics profiling postmortem^9^. The Seattle Alzheimer’s Disease Brain Cell Atlas (SEA-AD) provides quantitative, brain region-resolved neuropathology across multiple cortical and subcortical regions in donors from the Adult Changes in Thought study and the University of Washington Alzheimer’s Disease Research Center^24,25^. Combining these resources with matched dorsolateral prefrontal cortex proteomes and metabolomes from 391 donors, we show that AsymAD carries AD-level amyloid but fewer compact plaques, lower regional tau, and restricted TDP-43 pathology. A signature of preserved mitochondrial translation and respiration, synaptic proteins, polyunsaturated membrane lipids, and low immune signaling distinguishes AsymAD from AD within matched amyloid, tau, and TDP-43 strata, accompanied by pathology-specific adaptations. Within AsymAD, individuals who had developed MCI differ from those who remained cognitively intact in tangle burden and in episodic and semantic memory decline, yet retain a prefrontal proteome that is similar to that of controls and differs markedly in AD. These findings indicate that cognitive resilience reflects the preservation of a mitochondrial, synaptic, and lipid homeostatic state despite amyloid burden, and they identify processes that may need to be sustained to keep AD pathology clinically silent.

## Results

### Asymptomatic AD uncouples canonical AD pathology from cognitive decline and TDP-43 co-pathology

To define molecular features associated with resilience to Alzheimer’s disease (AD) pathology, we established a study framework spanning cognitively normal (controls; N = 85), asymptomatic AD (AsymAD; N = 169), and symptomatic AD (N = 137) individuals across the ROSMAP^23^ and SEA-AD^25^ cohorts (Figure 1A). Controls represented individuals with preserved cognition and limited AD neuropathology, whereas AsymAD individuals accumulated substantial amyloid-β and tau pathology without developing dementia. In contrast, individuals with AD exhibited both extensive neuropathology and cognitive impairment. We integrated detailed neuropathological measures with dorsolateral prefrontal cortex proteomic and metabolomic profiles and further examined individual pathologies across their regional stages of progression. Demographic and genetic characteristics, including age at death, sex, APOE genotype, and postmortem interval, were represented across diagnostic groups and cohorts (Figure 1B).

**Figure 1.**
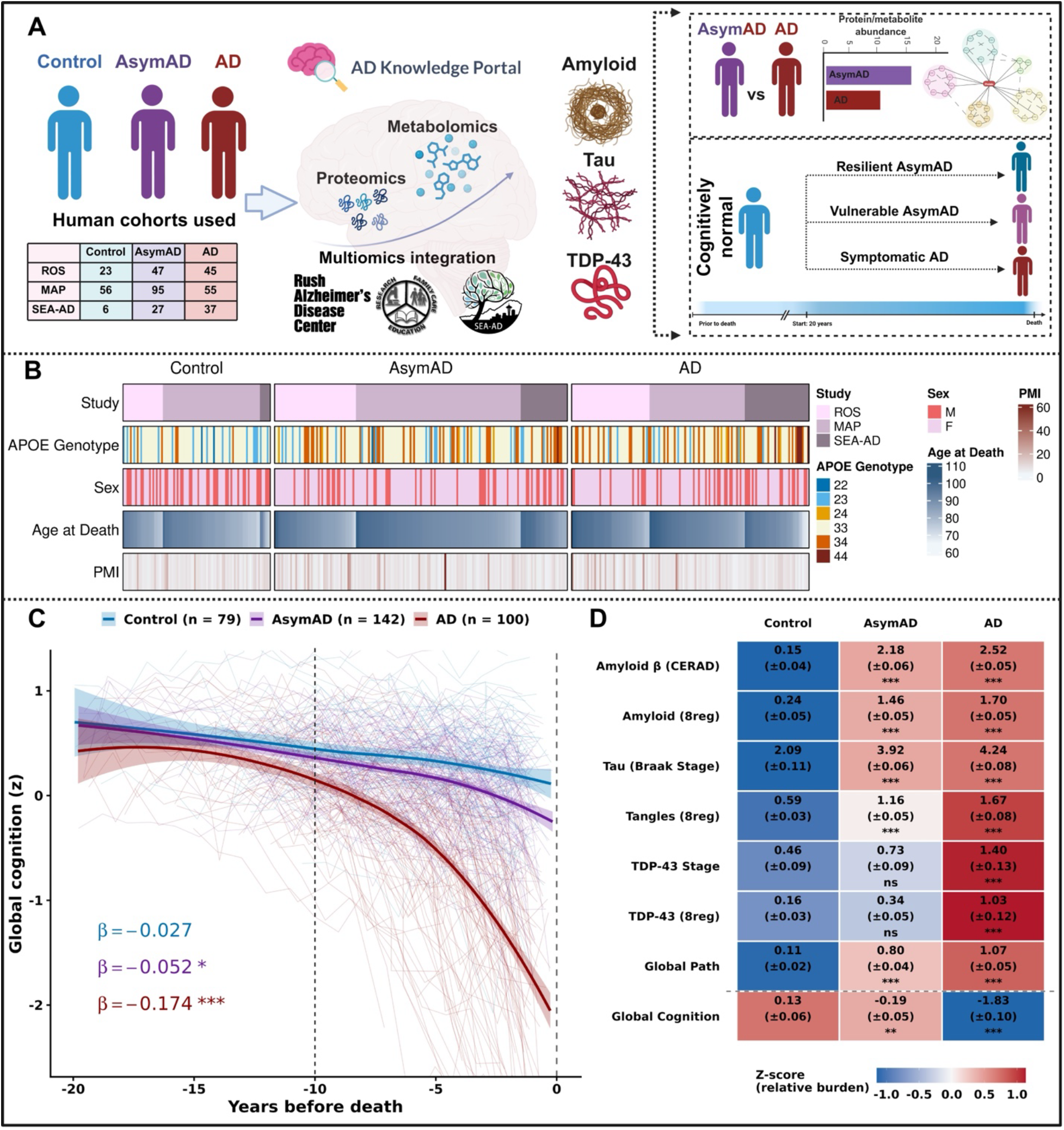
Multi-cohort design and clinicopathological characterization of control, AsymAD and AD donors. (A) Graphical representation of the study design. Postmortem brain donors from the Religious Orders Study and Rush Memory and Aging Project (ROSMAP; ROS and MAP) and the Seattle Alzheimer’s Disease Brain Cell Atlas (SEA-AD) were classified as control, asymptomatic AD (AsymAD) or symptomatic AD (AD). Donor numbers per cohort and group are shown in the table (ROS: 23/47/45; MAP: 56/95/55; SEA-AD: 6/27/37 for Control/AsymAD/AD). Bulk proteomic and metabolomic data from the AD Knowledge Portal were integrated across cohorts and related to amyloid, tau and TDP-43 pathology. Right, analytical framework: differential protein/metabolite abundance between AsymAD and AD, followed by stratification of pathology-burdened donors without dementia into resilient AsymAD, vulnerable AsymAD and symptomatic AD, relative to the ∼20-year window before death. (B) Donor-level annotation heatmaps for each diagnostic group showing study (ROS, MAP, SEA-AD), *APOE* genotype, sex (M, F), age at death and postmortem interval (PMI). Each column is one donor. (C) Longitudinal trajectories of global cognition (z-score) as a function of years before death in Control (blue, n = 79), AsymAD (purple, n = 142) and AD (red, n = 100) donors from ROSMAP. Thin lines show individual donor trajectories; thick lines show LOESS-smoothed group trajectories (span = 0.9) with shaded 95% confidence intervals. β, rate of cognitive change per year before death, estimated by a linear mixed-effects model with diagnosis × time, age at death (mean-centered), and a random intercept and slope per donor; asterisks indicate the p value (Satterthwaite t-test) for the difference between each group’s slope and Control’s (reference group, unmarked): *\*p < 0.05, **p < 0.01, ***p < 0.001*. (D) Neuropathological and cognitive profile of each group. Cell values are group mean (± s.e.m.) for CERAD score, Braak stage, TDP-43 stage, amyloid, tangle and TDP-43 burden [regional (“8reg”) composite], global pathology and global cognition. Color indicates the z-score of relative burden across groups (blue, lower; red, higher; scale at bottom). The dashed line separates pathology measures from cognition. Asterisks denote significance of each group versus Control by Dunn’s test, Benjamini-Hochberg correction: ns, not significant; \**p* < 0.05, \*\**p* < 0.01, \*\*\**p* < 0.001.

We first leveraged ROSMAP^23^ to examine longitudinal cognitive trajectories before death. Global cognition declined only modestly in controls (β = −0.027 SD/year) and AsymAD (β = −0.052 SD/year), whereas AD showed substantially steeper decline (β = −0.174 SD/year; Figure 1C). Separation between AsymAD and AD became most pronounced during the final decade of life. Relative preservation in AsymAD extended across all five cognitive domains, with AD declining more steeply than AsymAD in episodic memory, semantic memory, perceptual speed, perceptual orientation, and working memory (Figure S1A). AsymAD nonetheless showed modest but significant decline in episodic memory (β = −0.037 SD/year), semantic memory (β = −0.062 SD/year), and perceptual speed (β = −0.093 SD/year), but not in perceptual orientation or working memory. The timing of divergence also differed by domain. In AD, episodic and semantic memory began to separate from controls and AsymAD approximately 15 years before death, whereas working memory and perceptual orientation diverged only in the final 5–8 years. Perceptual speed was lower in AD throughout the observation window compared to AsymAD and controls (Figure S1A). The number of donors contributing data at each follow-up year is shown in Figure S1B.

Neuropathological profiling in ROSMAP revealed a marked dissociation between canonical AD pathology, TDP-43 co-pathology, and cognition (Figure 1D). AsymAD and AD both showed elevated amyloid and tau pathology relative to controls, although tangle burden and Braak stage were higher in AD, resulting in high global AD pathology in both groups (group means shown inside the box). In contrast, TDP-43 accumulation and spread remained substantially lower in AsymAD than in AD. Thus, despite carrying an AD-like amyloid and tau burden, AsymAD individuals maintained cognition while showing limited TDP-43 co-pathology.

We also investigated whether this phenotype was observed across brain regions in an independent cohort. In SEA-AD, AsymAD donors occupied CERAD and Braak stages overlapping those observed in AD despite the absence of dementia (Figure S2A). Region-resolved neuropathological profiling across 11 cortical, limbic, and subcortical regions reproduced the pattern observed in ROSMAP. AsymAD showed extensive amyloid and tau pathology across multiple regions, whereas TDP-43 burden remained limited and cognitive performance was relatively preserved compared to AD (Figure S2B; values in Table S1).

Together, these longitudinal and cross-cohort analyses identify AsymAD as a distinct pathological state in which substantial amyloid and tau accumulation is uncoupled from both widespread TDP-43 co-pathology and cognitive decline. The reproducibility of this phenotype across longitudinal cognitive assessment, independent cohorts, and multiple brain regions provided the foundation for investigating the molecular programs associated with resilience to AD pathology.

### Pathology composition distinguishes resilient AsymAD from symptomatic AD

To determine whether AsymAD and AD differ in the composition of pathology rather than its amount, we compared amyloid, tau, and TDP-43 burden across diagnostic groups and pathological stages. In ROSMAP, amyloid burden in AsymAD was substantially greater than in controls and comparable to AD within CERAD stages 2 and 3 (Figure 2A). Tau pathology followed a different pattern. AsymAD occupied an intermediate position between controls and AD overall, and tangle burden was significantly lower in AsymAD than in AD at Braak stages 4 and 5 but did not reach significance at stage 3. In contrast, TDP-43 burden in AsymAD did not differ from controls, whereas AD showed markedly greater TDP-43 accumulation, with a significant difference at TDP-43 stage 3.

**Figure 2.**
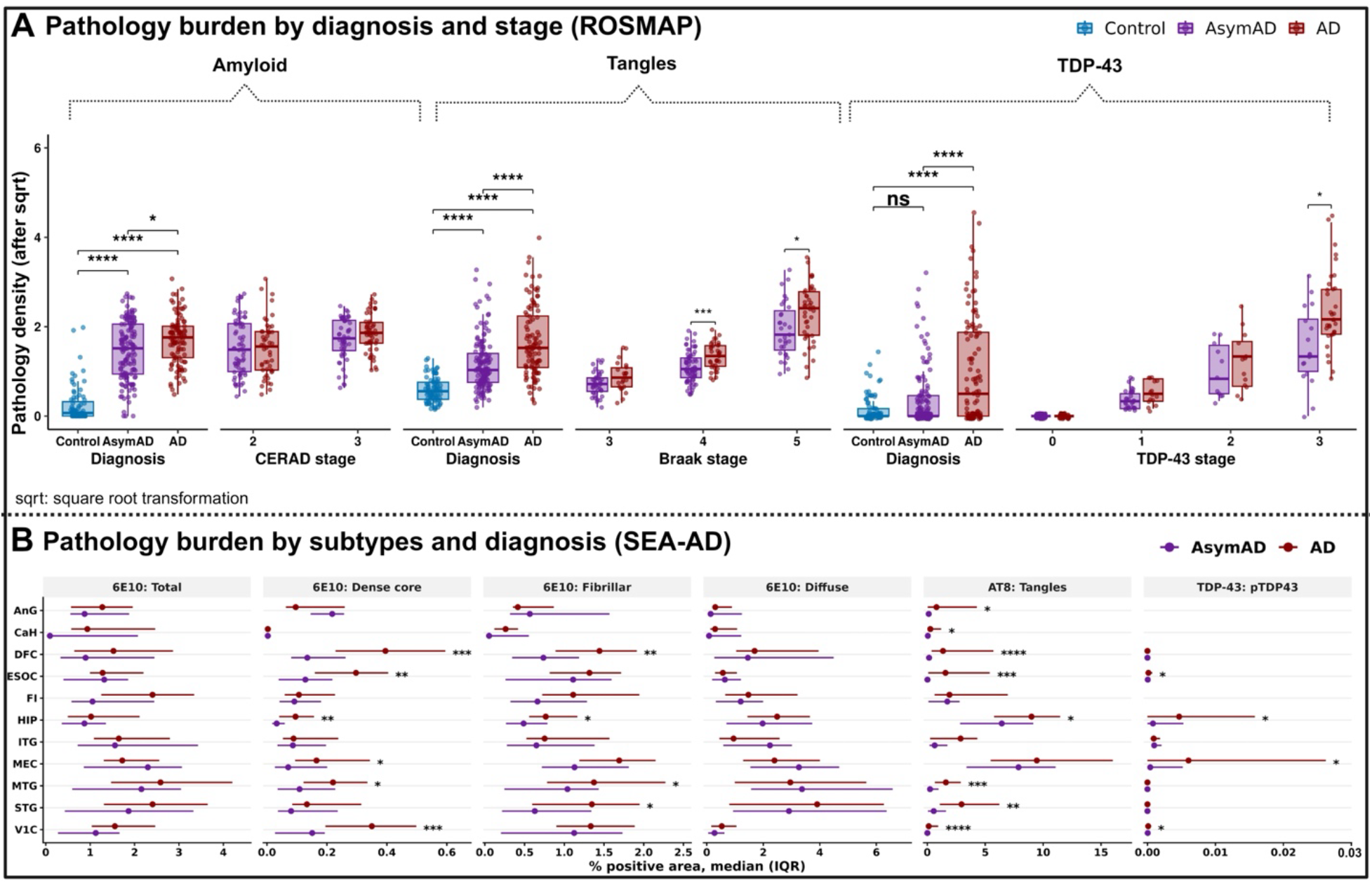
AsymAD donors carry AD-level amyloid but less tau and little TDP-43 pathology. (A) Pathology burden in ROSMAP by diagnosis and by neuropathological stage. Boxplots show amyloid, tangles and TDP-43 (8-region composite), each shown as square-root-transformed pathology density (y-axis) across Control (blue), AsymAD (purple) and AD (red). The stage panels compare AsymAD with AD within strata of CERAD stage (2, 3), Braak stage (3-5) and TDP-43 stage (0-3). Control donors are not shown in the stage panels. Points are individual donors; boxes show the median and interquartile range (IQR), and whiskers extend to 1.5 × IQR. Group sizes: Control (n = 79), AsymAD (n = 142), AD (n = 100). Brackets show pairwise comparisons by Dunn’s test with Benjamini–Hochberg adjustment: ns, not significant; \**p* < 0.05, \*\**p* < 0.01, \*\*\**p* < 0.001, \*\*\*\**p* < 0.0001. (B) Regional burden of amyloid subtypes, tangles and phospho-TDP-43 in SEA-AD AsymAD (purple, n = 27) and AD (red, n = 37) donors. Panels show 6E10-positive total amyloid, dense-core, fibrillar and diffuse plaques, AT8-positive tangles and pTDP43, quantified as percent positive area in angular gyrus (AnG), head of caudate (CaH), dorsolateral prefrontal cortex (DFC), extrastriate occipital cortex (ESOC), frontoinsula (FI), hippocampus (HIP), inferior temporal gyrus (ITG), medial entorhinal cortex (MEC), middle temporal gyrus (MTG), superior temporal gyrus (STG) and primary visual cortex (V1C). Points show the median and lines the IQR. pTDP43 was not quantified in AnG and CaH. Asterisks (right of each region) indicate AsymAD versus AD differences by two-sided Wilcoxon rank-sum test, Benjamini–Hochberg-adjusted across regions within each panel; only significant differences are marked: *\*p < 0.05, **p < 0.01, ***p < 0.001, ****p < 0.0001*.

To test whether the lower TDP-43 burden in AsymAD might be related to its comparatively lower tau burden despite extensive amyloid pathology, we examined pairwise associations among amyloid, tangles, and TDP-43 (Figure S3A). Amyloid burden was positively associated with tangle burden in both AsymAD (slope = 0.346) and AD (slope = 0.516), but not in controls (slope = −0.087). Tangle burden was also positively associated with TDP-43 pathology in AsymAD (slope = 0.202) and AD (slope = 0.379), whereas this relationship was absent in controls (slope = 0.082). In contrast, TDP-43 showed little relationship with amyloid burden in AsymAD (slope = −0.099) and only a weak positive relationship in AD (slope = 0.104; control slope = −0.007). These findings suggest that TDP-43 accumulation is more closely associated with tau burden than with amyloid burden alone, consistent with the lower tau and TDP-43 pathology observed in AsymAD despite substantial amyloid accumulation.

Because ROSMAP pathology burden represents a composite measure across eight brain regions^26^, we used SEA-AD to determine whether pathological differences between AsymAD and AD were region- and amyloid-subtype specific. Total amyloid burden was broadly comparable between AsymAD and AD across the 11 examined regions (Figure 2B), with relatively low accumulation in the head of caudate (CaH) and greater burden in regions including the frontoinsula (FI) and middle temporal gyrus (MTG). However, decomposition of total amyloid into pathological subtypes uncovered differences that were obscured at the level of total burden.

Dense-core amyloid was greater in AD than AsymAD in several regions, including the dorsolateral prefrontal cortex (DFC), extrastriate occipital cortex (ESOC), hippocampus (HIP), medial entorhinal cortex (MEC), middle temporal gyrus (MTG), and primary visual cortex (V1C). Fibrillar amyloid was similarly increased in AD in selected regions, including the dorsolateral prefrontal cortex (DFC), hippocampus (HIP), middle temporal gyrus (MTG), and superior temporal gyrus (STG). In contrast, diffuse amyloid showed substantially greater overlap between AsymAD and AD. Thus, similar total amyloid burden can conceal important differences in pathological composition, with AD showing greater enrichment of compact amyloid species across vulnerable regions compared to AsymAD.

Tau pathology further distinguished pathological states in AsymAD and AD. AT8-positive tangle burden was greater in AD across several regions, including the dorsolateral prefrontal cortex (DFC), extrastriate occipital cortex (ESOC), middle temporal gyrus (MTG), superior temporal gyrus (STG), and primary visual cortex (V1C) (Figure 2B). Regional pTDP-43 showed an even more restricted distribution, with the clearest increases in AD observed in the hippocampus (HIP) and medial entorhinal cortex (MEC). Analysis across all 11 SEA-AD regions confirmed that total amyloid burden was highly similar between AsymAD and AD across all CERAD stages; however amyloid subtype-specific differences were detected in distinct brain regions, including DFC, ESOC, V1C, and HIP (Figure S3B). AT8-positive tau burden increased with advancing Braak stage, with regional accumulation being higher in AD than in AsymAD (Figure S3C). Regional pathology values are provided in Table S2.

These analyses indicate that while AsymAD involves widespread amyloid and significant tau accumulation, it features distinct pathological composition and spatial distribution. Despite a consistently high overall amyloid burden, vulnerable brain regions show lower enrichment of dense core and fibrillar amyloid species. We also observed that regional tau accumulation is lower, and progression toward extensive TDP-43 co-pathology is severely limited.

### AsymAD exhibits coordinated metabolic remodeling of the prefrontal cortex

To define the molecular programs associated with resilience, we profiled the proteome and metabolome of the dorsolateral prefrontal cortex (DFC; BA9), a region central to executive function that is progressively affected in AD^25^. Proteomic profiles were available from 242 donors (142 AsymAD, 100 AD) and metabolomic profiles from 326 donors (147 AsymAD, 179 AD), of whom 166 had matched measurements across both modalities (93 AsymAD, 73 AD; Figure 3A). After adjustment for age at death, sex, postmortem interval, and APOE genotype, 799 proteins and 53 metabolites differed between AsymAD and AD (adjusted p < 0.05). Of these, 474 proteins and 27 metabolites were more abundant in AsymAD, and 325 proteins and 26 metabolites were less abundant in AsymAD compared to AD (Figures 3A-3C).

**Figure 3.**
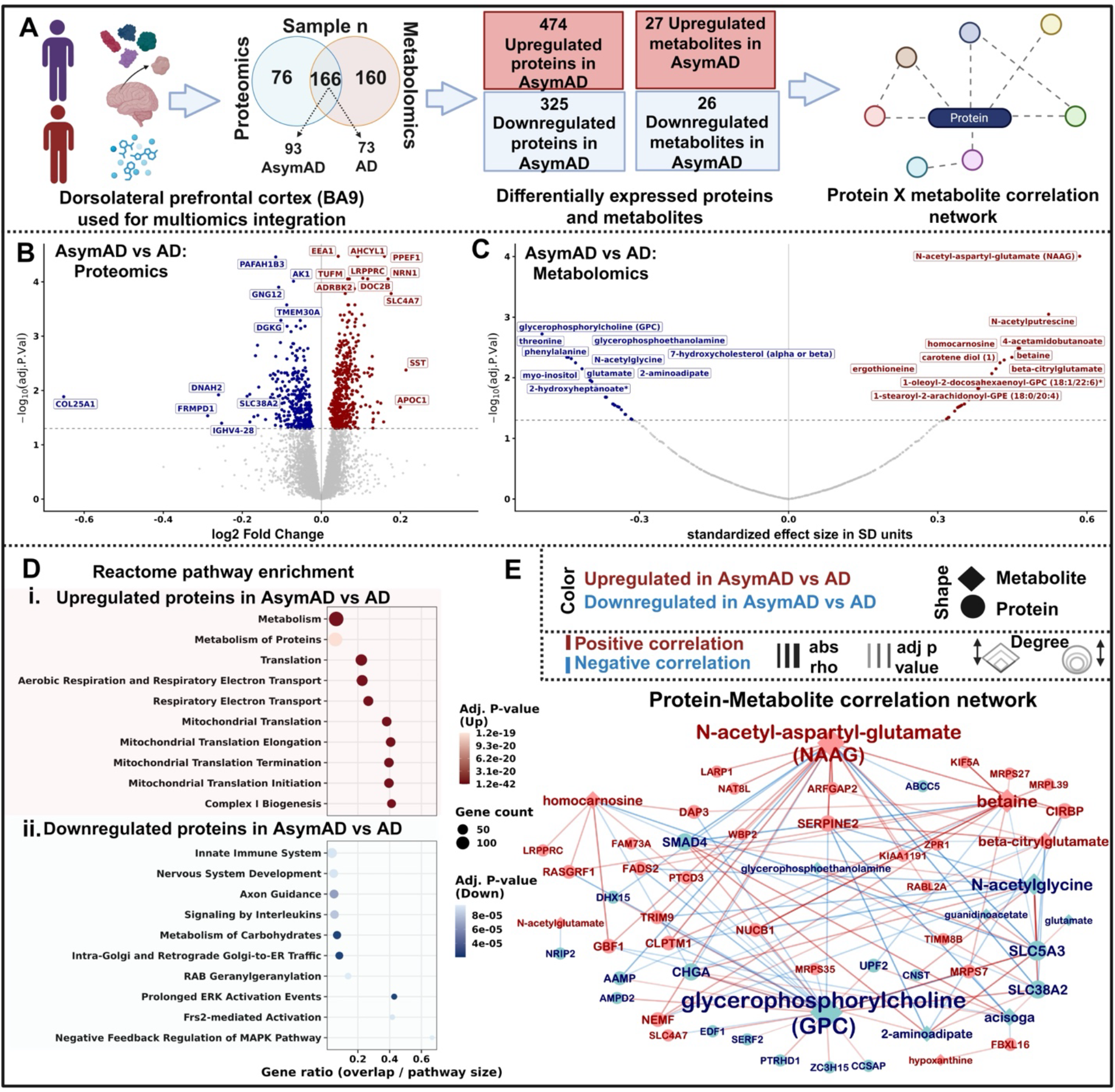
AsymAD is characterized by coordinated mitochondrial, neuronal, and metabolic remodeling of the dorsolateral prefrontal cortex. (A) Study design. Proteomic (n = 242; 142 AsymAD, 100 AD) and metabolomic (n = 326; 147 AsymAD, 179 AD) profiles were generated from the dorsolateral prefrontal cortex (BA9), with 166 donors (93 AsymAD, 73 AD) having matched measurements across both modalities. Differential abundance analysis identified 474 proteins and 27 metabolites higher, and 325 proteins and 26 metabolites lower, in AsymAD than in AD. Matched samples were used to construct a protein–metabolite correlation network. (B and C) Volcano plots of differentially regulated proteins (B) and metabolites (C) abundance between AsymAD and AD. The x axis shows log2 fold change (B) or standardized effect size in SD units (C), with positive values indicating higher abundance in AsymAD, and the y axis shows −log10 adjusted p value. Red indicates features higher in AsymAD, blue indicates features higher in AD, and gray indicates non-significant features. The dashed line marks adjusted p = 0.05. Selected top-ranked features are labeled. Metabolites marked with an asterisk denote compounds identified with lower confidence. (D) Reactome pathway enrichment of proteins higher (i) or lower (ii) in AsymAD. Dot position indicates gene ratio (overlap/pathway size), dot size indicates the number of overlapping proteins, and dot color indicates adjusted p value. (E) Protein–metabolite correlation network constructed from the 166 donors with matched proteomic and metabolomic profiles. Diamonds represent metabolites and circles represent proteins. Node color indicates direction of change (red, higher in AsymAD; blue, lower in AsymAD), and node size is proportional to degree. Edge color indicates positive (red) or negative (blue) correlation, edge width is proportional to absolute Spearman’s ρ, and edge opacity reflects adjusted p value. Only correlations with |ρ| ≥ 0.5 and adjusted p < 0.05 are shown. Differential abundance was assessed with limma, adjusting for age at death, sex, postmortem interval, and APOE genotype. p values were adjusted by the Benjamini-Hochberg method.

The proteins most significantly more abundant in AsymAD than in AD converged on endosomal trafficking (EEA1), mitochondrial translation (TUFM, LRPPRC), and synaptic function (NRN1, DOC2B, SST) (Figure 3B). Proteins lower in abundance in AsymAD pointed to differences in membrane lipid handling and cellular energy buffering. They included the phospholipid-modifying enzymes PAFAH1B3 and DGKG, TMEM30A, the β-subunit of the flippase complexes that maintain phospholipid asymmetry, the adenylate kinase AK1, and the amino acid transporter SLC38A2. All differentially abundant proteins are listed in Table S3. Pathway enrichment placed these changes within a coherent bioenergetic program. Proteins elevated in AsymAD were strongly enriched for cytosolic and mitochondrial translation, aerobic respiration and respiratory electron transport, and complex I biogenesis (Figure 3Di). Proteins reduced in AsymAD were enriched for innate immune and interleukin signaling, carbohydrate metabolism, Golgi-to-ER trafficking, RAB geranylgeranylation, and MAPK/ERK signaling cascades (Figure 3Dii). Resilience was therefore associated with preserved mitochondrial translation and respiratory capacity, alongside attenuated inflammatory and growth-factor signaling.

The metabolome independently supported preserved neuronal function and reduced membrane stress in AsymAD. N-acetylaspartylglutamate (NAAG), the most abundant peptide neurotransmitter in the brain and a neuron-specific modulator of glutamatergic transmission^27^, was the most significantly elevated metabolite in AsymAD. It was accompanied by its biosynthetic relative beta-citrylglutamate, the brain-enriched antioxidant dipeptide homocarnosine, the methyl donor betaine, and the antioxidant ergothioneine (Figure 3C). AsymAD brains also had higher levels of intact membrane phospholipids carrying polyunsaturated fatty acids (PUFAs), including PC(18:1/22:6) and PE(18:0/20:4). In contrast, glycerophosphorylcholine (GPC) and glycerophosphoethanolamine, which are breakdown products of these phospholipids^28^, were lower in AsymAD alongside glutamate, myo-inositol, and the oxysterol 7-hydroxycholesterol (Figure 3C). All differentially abundant metabolites are listed in Table S3. In AsymAD, higher NAAG with lower glutamate is consistent with restrained excitatory signaling^29,30^, and higher intact phospholipids with fewer breakdown products are consistent with less membrane breakdown^31^.

To determine how the differentially abundant protein and metabolite changes are coordinated between AsymAD and AD, we constructed a correlation network from the 166 donors with matched profiles (Figure 3E). The two most connected metabolite hubs were NAAG, which was higher in AsymAD than in AD, and GPC, which was lower. NAAG had 20 protein neighbors of which 13 correlated positively and were all higher in AsymAD, including the trafficking protein ARFGAP2, the neuronal enzyme NAT8L, and the matrix protein SERPINE2. The 7 that correlated negatively were all lower in AsymAD, including the transporter SLC5A3 and the signaling protein SMAD4. GPC had 26 neighbors: 13 proteins lower in AsymAD, such as the transporter SLC38A2 and the RNA factor UPF2, correlated positively with GPC. Thirteen higher in AsymAD, including the mitochondrial proteins DAP3 and MRPS7, the neuronal protein RASGRF1, and the lipid enzyme FADS2, correlated with GPC negatively. Betaine, homocarnosine, and beta-citrylglutamate, which are higher in AsymAD, correlated with the shared proteins with the same sign as NAAG and the opposite sign from GPC. The axonal protein KIF5A and the mitochondrial proteins MRPS27 and MRPL39 correlated positively with betaine, and the mitochondrial protein LRPPRC with homocarnosine.

Two subnetworks drawn from the same correlation data matched processes highlighted in Figures 3B–3D (Figure S4; thresholds and module definitions are in the Methods). The first contained 34 mitochondrial proteins, all higher in AsymAD, consistent with the mitochondrial translation enrichment in Figure 3D. Most were mitoribosomal subunits, and the rest included LRPPRC, SLIRP, and the import and assembly factors TIMM8B, TRMT10C, and NDUFAF7. Their correlations with homocarnosine and betaine were all positive, and their correlations with 2-aminoadipate (a lysine-degradation intermediate and marker of protein oxidation) and acisoga (a polyamine oxidation product) were all negative (Figure S4A). The second module centered on membrane lipids. PUFA-containing phospholipids correlated with 50 proteins (Figure S4B), including the synaptic proteins CACNG3 and RPH3A, the fatty acid desaturase FADS2, mitochondrial translation proteins, and ATP8A1 and TMEM30A, two phospholipid flippase subunits that are lower in AsymAD. Thirty-two of these proteins also correlated with GPC or glycerophosphoethanolamine, and for each of them the sign for the intact phospholipids was opposite to the sign for their breakdown products (Table S3).

Together, these analyses indicate that there are distinct protein-metabolite interactions in AsymAD that govern the physiological changes. Mitochondrial translation is coupled to neuronal metabolites and intact polyunsaturated membrane lipids, and inversely to oxidative catabolites and phospholipid breakdown products.

### Stratification by proteinopathy stage reveals a conserved mitochondrial core and pathology-specific adaptations in AsymAD

To investigate how AsymAD brains maintain cognitive function despite different proteinopathies, we compared the DLPFC (BA9) proteome of AsymAD and AD across different stages of amyloid, tau, and TDP-43 pathology. The low and high stages were moderate (CERAD 2; 66 AsymAD, 48 AD) and frequent (CERAD 3; 51 AsymAD, 52 AD) neuritic plaque density for amyloid; limbic (Braak III–IV; 111 AsymAD, 57 AD) and neocortical (Braak V; 31 AsymAD, 40 AD) involvement for tau; and pathology confined to the amygdala, entorhinal cortex, and hippocampus (LATE-NC stages 1–2; 42 AsymAD, 28 AD) or extending to the neocortex (LATE-NC stage 3; 17 AsymAD, 33 AD) for TDP-43. Each proteinopathy was associated with hundreds of stratum-specific differences, in both directions (Figure 4A). At the low stage, 104 proteins were more abundant and 93 less abundant in AsymAD than in AD under amyloid, 304 and 349 under tau, and 217 and 170 under TDP-43. At the high stage, the amyloid-specific signature increased about five-fold, to 454 more abundant and 570 less abundant proteins, despite similar donor numbers, whereas the tau-specific (116 and 169) and TDP-43-specific (150 and 162) signatures were smaller, although their AsymAD groups were also smaller (31 and 17 donors). Thus, amyloid stage had the largest effect on the AsymAD vs AD proteomic difference. All differentially abundant proteins are listed in Table S4.

**Figure 4.**
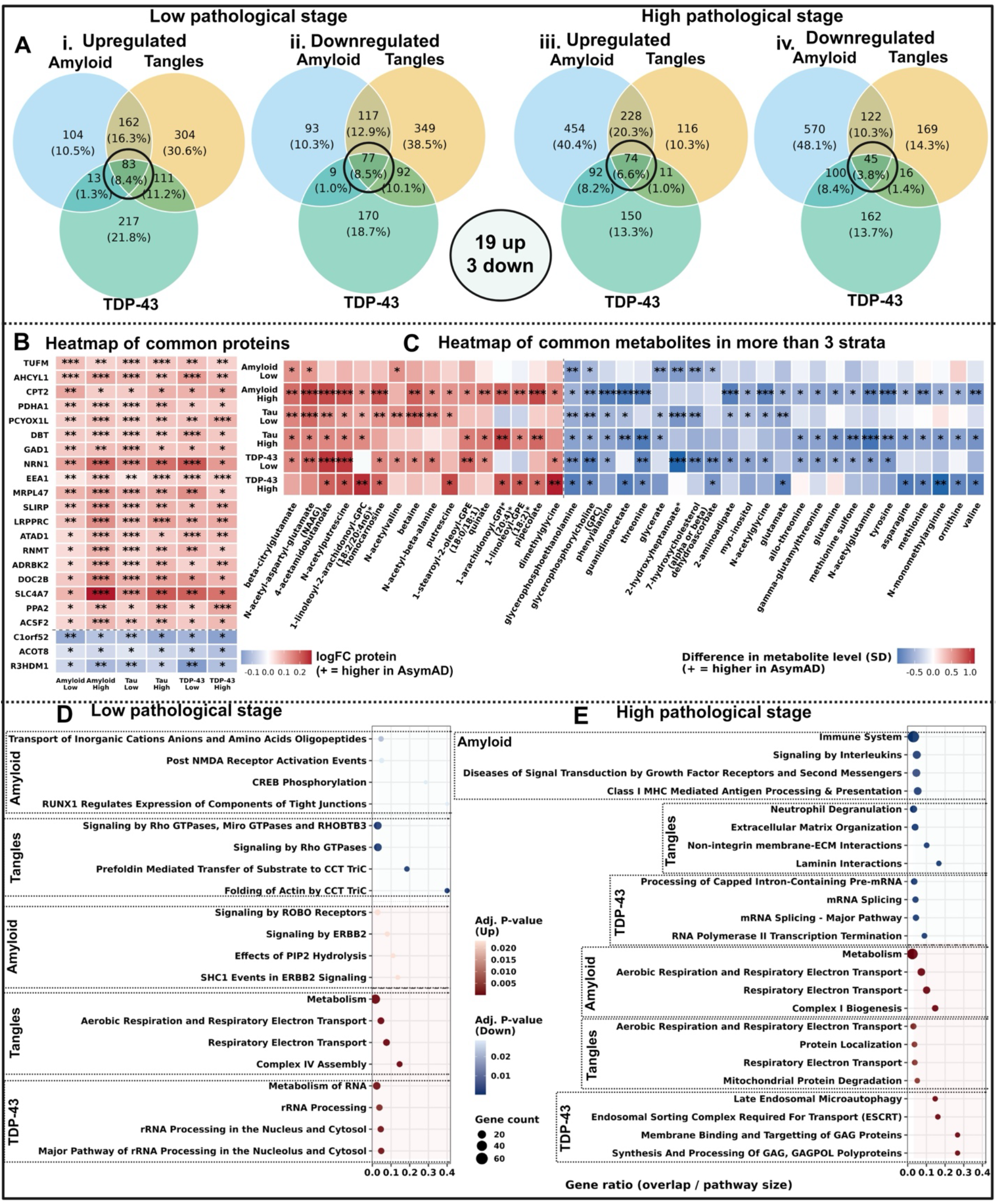
Stratification by proteinopathy stage reveals a conserved mitochondrial core and pathology-specific adaptations in AsymAD. (A) Venn diagrams of proteins differentially abundant between AsymAD and AD in the DFC (BA9) within amyloid, tau, and TDP-43 strata. Proteins higher (i, iii) and lower (ii, iv) in AsymAD are shown for the low (i, ii) and high (iii, iv) pathological stages. Values are the number of proteins in each region, with the percentage of all proteins in the diagram in parentheses. Low-stage strata were moderate amyloid (CERAD 2; n = 66 AsymAD, 48 AD), limbic tau (Braak III–IV; n = 111, 57), and early TDP-43 (LATE-NC stages 1–2; n = 42, 28). High-stage strata were frequent amyloid (CERAD 3; n = 51, 52), neocortical tau (Braak V; n = 31, 40), and neocortical TDP-43 (LATE-NC stage 3; n = 17, 33). Black circles mark proteins altered in all three proteinopathies at each stage. Nineteen proteins were higher and 3 lower in AsymAD across all six strata. (B) Heatmap of log2 fold change (AsymAD versus AD) for the 22 proteins shared across all strata. Columns show each proteinopathy at the low and high stage. Red indicates higher and blue lower abundance in AsymAD. *\*p < 0.05, **p < 0.01, ***p < 0.001 (nominal)*. (C) Heatmap of standardized effect size (SD units; AsymAD versus AD) for the 40 metabolites differing in the same direction in at least three of the six strata: 16 higher (left) and 24 lower (right of the dashed line) in AsymAD. Asterisks as in (B). Metabolite names marked with an asterisk denote compounds identified with lower confidence. (D and E) Reactome pathway enrichment of proteins uniquely altered within each proteinopathy at the low (D) and high (E) pathological stage. Pathways are grouped by pathology; pink shading marks pathways enriched among proteins higher in AsymAD and blue shading those enriched among proteins lower in AsymAD. Dot position indicates gene ratio (overlap/pathway size), dot size the number of overlapping proteins, and dot color adjusted p value.

Alongside these stratum-specific changes, 22 proteins distinguished AsymAD from AD in all six strata, 19 more abundant and 3 less abundant in AsymAD (Figures 4A and 4B). The more abundant core was dominated by mitochondrial proteins governing translation (TUFM, MRPL47, LRPPRC, SLIRP), pyruvate, branched-chain amino acid, and fatty acid metabolism (PDHA1, DBT, CPT2, ACSF2), and mitochondrial homeostasis (ATAD1, PPA2). It also included proteins supporting inhibitory neurotransmission and synaptic function (GAD1, NRN1, DOC2B) and the early endosomal protein EEA1, which was among the most significantly elevated proteins in the unstratified comparison (Figure 3B). Only C1orf52, ACOT8, and R3HDM1 were consistently less abundant in AsymAD. Mitochondrial and neuronal maintenance therefore emerges as a stable feature of AsymAD that does not depend on the type or stage of pathology.

To identify programs engaged by each proteinopathy, we performed pathway enrichment on the uniquely altered proteins (Figures 4D and 4E). At the low stage, proteins uniquely lower in AsymAD under amyloid were enriched for inorganic ion and amino acid transport, post-NMDA receptor activation events, and CREB phosphorylation, suggesting dampened excitatory signaling. Under tau, less abundant proteins were enriched for Rho GTPase signaling and CCT/TRiC-mediated cytoskeletal protein folding. TDP-43-specific proteins less abundant in AsymAD yielded no significant pathways. Proteins uniquely higher in AsymAD were enriched for ROBO and ERBB2 signaling under amyloid, aerobic respiration, electron transport, and complex IV assembly under tau, and RNA metabolism and rRNA processing under TDP-43 (Figure 4D).

At the high stage, these programs diverged further (Figure 4E). Under frequent amyloid, less abundant proteins were enriched for immune system, interleukin signaling, growth-factor receptor signaling, and class I MHC antigen presentation, whereas more abundant proteins were enriched for metabolism, respiratory electron transport, and complex I biogenesis. Under neocortical tau, less abundant proteins were enriched for neutrophil degranulation and extracellular matrix organization, including laminin and non-integrin membrane–ECM interactions, and more abundant proteins for aerobic respiration, protein localization, and mitochondrial protein degradation. Under neocortical TDP-43, less abundant proteins were enriched for pre-mRNA processing and splicing, and more abundant proteins for late endosomal microautophagy and ESCRT-mediated membrane trafficking. Thus, as pathology advances, AsymAD couples sustained respiratory capacity with suppressed immune and matrix-remodeling responses under amyloid and tau, and with altered RNA processing and enhanced endolysosomal clearance under TDP-43.

To determine whether these adaptations were mirrored in the metabolome, we applied the same stratification to the BA9 metabolomic data (Figure S5). Differences were again both shared and pathology-specific: of the 66 metabolites higher in AsymAD at the low stage, 25 were specific to tau and 23 to TDP-43, whereas at the high stage 31 of 86 were specific to amyloid. Forty metabolites differed in the same direction in at least three strata (Figure 4C). Sixteen were higher in AsymAD, including the neuronal metabolites NAAG, beta-citrylglutamate, and homocarnosine, the methyl donors betaine and dimethylglycine, and polyunsaturated phospholipids carrying arachidonate or docosahexaenoate. Twenty-four were lower, including the phospholipid breakdown products GPC and glycerophosphoethanolamine, the lysine-degradation intermediate 2-aminoadipate, the oxysterol 7-hydroxycholesterol, the astrocyte-associated metabolite myo-inositol, and glutamate, together with several amino acids such as tyrosine, phenylalanine, and valine. All differentially abundant metabolites are listed in Table S4.

Across all six strata, 22 proteins, mostly mitochondrial, synaptic, and endosomal, distinguished AsymAD from AD. Each proteinopathy also had its own changes: higher respiration under amyloid and tau, lower immune signaling under amyloid, lower matrix remodeling under tau, and higher endolysosomal clearance under TDP-43.

### Higher tau pathology separates vulnerable from resilient AsymAD

We classified donors by combining longitudinal clinical diagnoses, collected up to 15 years before death, with autopsy pathology (Figure 5A). Controls (n = 79) had no dementia and less AD pathology. Three groups had AD-level pathology: resilient AsymAD (n = 77), who remained cognitively unimpaired (NCI) until death; vulnerable AsymAD (n = 65), who had mild cognitive impairment (MCI) without dementia; and AD (n = 100), who had dementia. More than half of the donors who were cognitively unimpaired at their final evaluation (77 of 144) had AD-level pathology at autopsy, so clinical status alone would have grouped them with controls. Ordering the groups from controls through resilient and vulnerable AsymAD to AD, we found that resilient and vulnerable AsymAD differed in tangle burden and cognition but not in amyloid or TDP-43 pathology (Figure 5B). Tangles increased at every step of this order, whereas amyloid was elevated relative to controls but did not differ among resilient AsymAD, vulnerable AsymAD, and AD, and TDP-43 increased only in AD. Global cognition declined more than twice as fast in vulnerable as in resilient AsymAD (β = −0.072 vs. −0.033 SD/year; Figure S6A). This difference was confined to episodic memory (β = −0.070 vs. −0.007 SD/year) and semantic memory (β = −0.086 vs. −0.040 SD/year) and was absent for working memory, perceptual speed, and perceptual orientation. Thus, among individuals with comparable amyloid and TDP-43 burden, vulnerability coincided with greater tangle burden and selective loss of the memory domains that depend on the temporal lobe, where tangles first accumulate.

**Figure 5.**
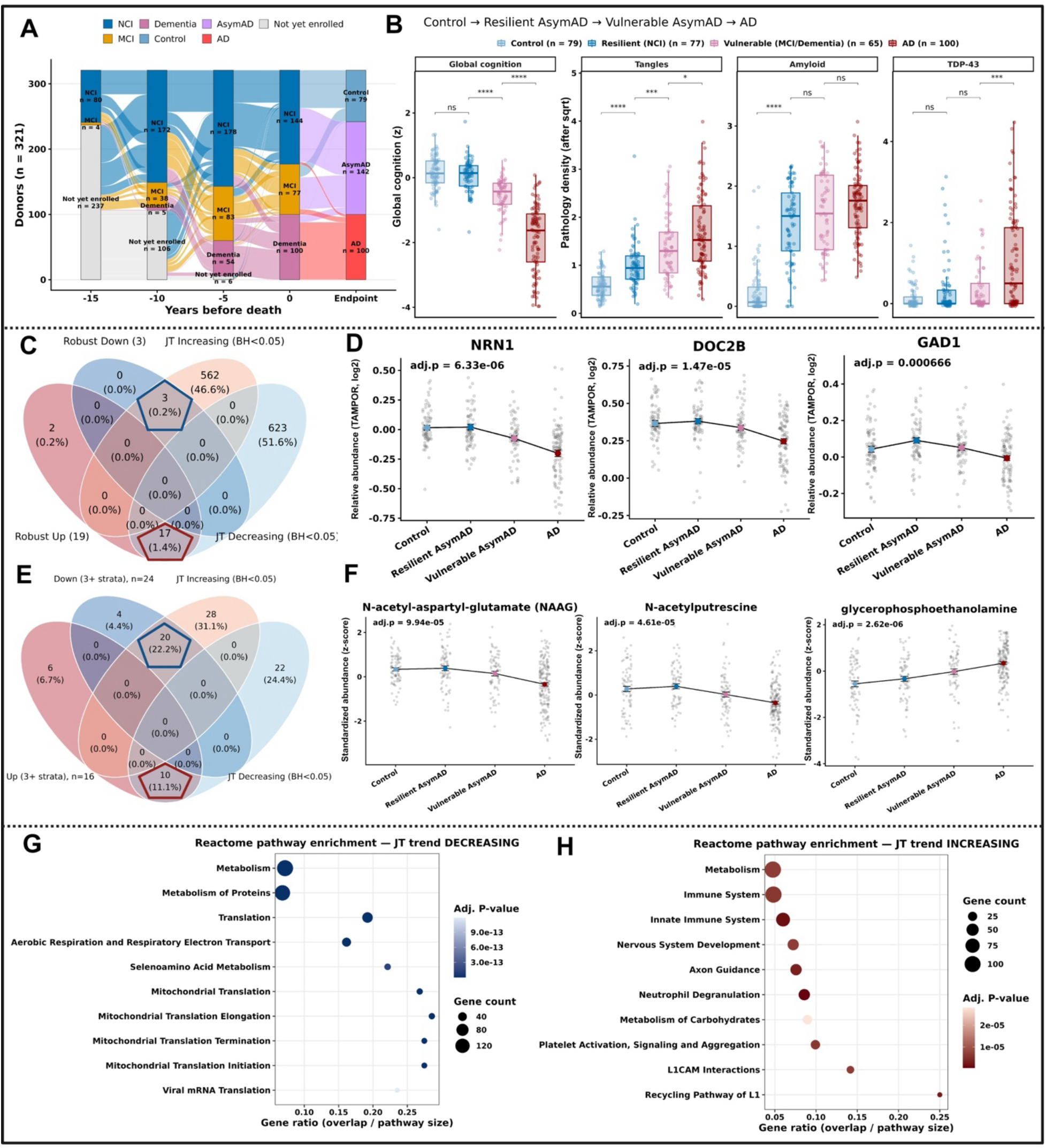
Vulnerability in AsymAD is linked to greater tangle accumulation. (A) Alluvial plot of clinical diagnoses at 15, 10, 5, and 0 years before death for 321 donors (NCI, no cognitive impairment; MCI, mild cognitive impairment), with the final group at the endpoint. (B) Global cognition and neuropathology (tangles, amyloid, TDP-43) in controls (n = 79), resilient AsymAD (NCI; n = 77), vulnerable AsymAD (MCI; n = 65), and AD (n = 100). Boxes show median and interquartile range, with whiskers at 1.5× the interquartile range. Adjacent groups were compared by Wilcoxon rank-sum test with Benjamini–Hochberg correction: ns, not significant; *p < 0.05; ***p < 0.001; ****p < 0.0001. (C) Overlap between the stage-invariant AsymAD protein signature (Figure 4B) and proteins with significant Jonckheere–Terpstra trends (adjusted p < 0.05). (D) NRN1, DOC2B, and GAD1 (TAMPOR-normalized log2 abundance). (E) Overlap between metabolites higher or lower in AsymAD in at least three pathological strata (Figure 4) and metabolites with significant trends. (F) NAAG, N-acetylputrescine, and glycerophosphoethanolamine (z-scored abundance). In (D) and (F), colored points and error bars show mean ± SEM, gray points show individual donors, and adjusted p values are from the Jonckheere–Terpstra test with Benjamini–Hochberg correction. (G and H) Reactome enrichment of proteins decreasing (G) or increasing (H) along the continuum; dot position shows gene ratio, size the number of overlapping proteins, and color the adjusted p value.

We also compared each pair of groups across all 8,339 quantified BA9 proteins using limma^32^ (Figure S7A; Table S5). AD differed markedly from the other groups, with 889 proteins differing from controls and 516 from resilient AsymAD. The proteins most strongly lower in AD than in resilient AsymAD included the synaptic proteins NRN1, RPH3A, DOC2B, and GAD1, whereas PAFAH1B3, DGKG, and TMEM30A were higher. In contrast, AsymAD differed little from controls. Only 10 proteins distinguished resilient AsymAD and 19 distinguished vulnerable AsymAD, almost all higher and related to amyloid deposition or the extracellular matrix (MDK, SMOC1, APP, CTHRC1, and NTN1). No protein differed between resilient and vulnerable AsymAD, and only five differed between vulnerable AsymAD and AD, three of them mitochondrial proteins lower in AD (LRPPRC, TUFM, and SLC25A46).

To test for graded changes along the continuum, we applied a Jonckheere–Terpstra trend test, which identified 640 decreasing and 565 increasing proteins (adjusted p < 0.05; Figure 5C). Consistent with the pairwise results, these trends were driven largely by the AD group. The stage-invariant AsymAD signature from Figure 4B was concordant: 17 of 19 elevated proteins decreased toward AD, and all three reduced proteins increased, whereas CPT2 and ACSF2 showed no monotonic trend (adjusted p = 0.34 and 0.20). NRN1, DOC2B, and GAD1 were maintained at control levels in resilient AsymAD and lowest in AD, although differences between the two AsymAD subgroups did not survive correction (Figure 5D). Metabolites behaved similarly: of the 16 metabolites more abundant in AsymAD in at least three strata, 10 decreased toward AD, and of the 24 less abundant, 20 increased (Figure 5E). NAAG followed the same course, whereas glycerophosphoethanolamine increased across all four groups (Figure 5F).

Pathway enrichment of the trending proteins showed that proteins decreasing along the continuum were enriched for translation, mitochondrial translation, and aerobic respiration, whereas proteins increasing were enriched for innate immune signaling, neutrophil degranulation, carbohydrate metabolism, axon guidance, and L1CAM interactions (Figures 5G and 5H).

To evaluate individual proteins behavior across groups, we selected representative trending proteins annotated to mitochondrial, synaptic, and immune pathways (Figure S7B). Mitoribosomal subunits, complex I proteins (NDUFB7, NDUFB9, NDUFAF7), PDHA1, and IDH2, together with synaptic proteins including SNAP25, STX1A, CPLX1, SYT12, GRIA3, and GABRG2, were lowest in AD. Immune proteins including ICAM1, ITGAX, MYD88, and RAB27A were highest in AD. In all three modules, resilient and vulnerable AsymAD resembled controls. Representative metabolites showed the same pattern: the phospholipid breakdown products GPC and glycerophosphoethanolamine, 7-hydroxycholesterol, and 2-hydroxyheptanoate were higher in AD, whereas intact phospholipids, NAAG, beta-citrylglutamate, homocarnosine, and betaine were lower, and none differed significantly from controls in either AsymAD subgroup. Thus, the mitochondrial, synaptic, and immune alterations that distinguish AsymAD from AD are largely confined to AD, and vulnerable AsymAD retains a control-like bulk prefrontal proteome despite greater tangle burden and memory decline.

## Discussion

In this study, we performed systematic analyses to identify the pathological and molecular features that distinguish cognitively resilient individuals from those with symptomatic AD. By integrating longitudinal cognitive scores, regional neuropathology, and matched prefrontal proteomes and metabolomes from 391 donors across ROSMAP^23^ and SEA-AD^25^, we found that AsymAD individuals carry AD-level amyloid burden but exhibit fewer dense-core and fibrillar plaques, lower regional tau, and restricted TDP-43 pathology. At the molecular level, AsymAD was characterized by higher levels of mitochondrial translation, respiration, synaptic, and endolysosomal proteins, together with lower immune signaling and higher neuronal and polyunsaturated phospholipid metabolites. This molecular signature persisted across matched amyloid, tau, and TDP-43 pathology strata. Stratification by individual proteinopathies further revealed distinct molecular adaptations associated with amyloid, tau, and TDP-43 pathology, involving immune signaling, extracellular matrix organization and respiration, and endolysosomal clearance, respectively (Figure 6). Within the AsymAD group, vulnerable individuals differed from resilient individuals in tangle burden and in the trajectories of episodic and semantic memory decline, but not in amyloid or TDP-43 pathology. Despite these clinical and pathological differences, the prefrontal proteomes of both AsymAD groups largely resembled those of controls, apart from amyloid-associated proteins, whereas mitochondrial, synaptic, and immune alterations were predominantly observed in AD.

**Figure 6.**
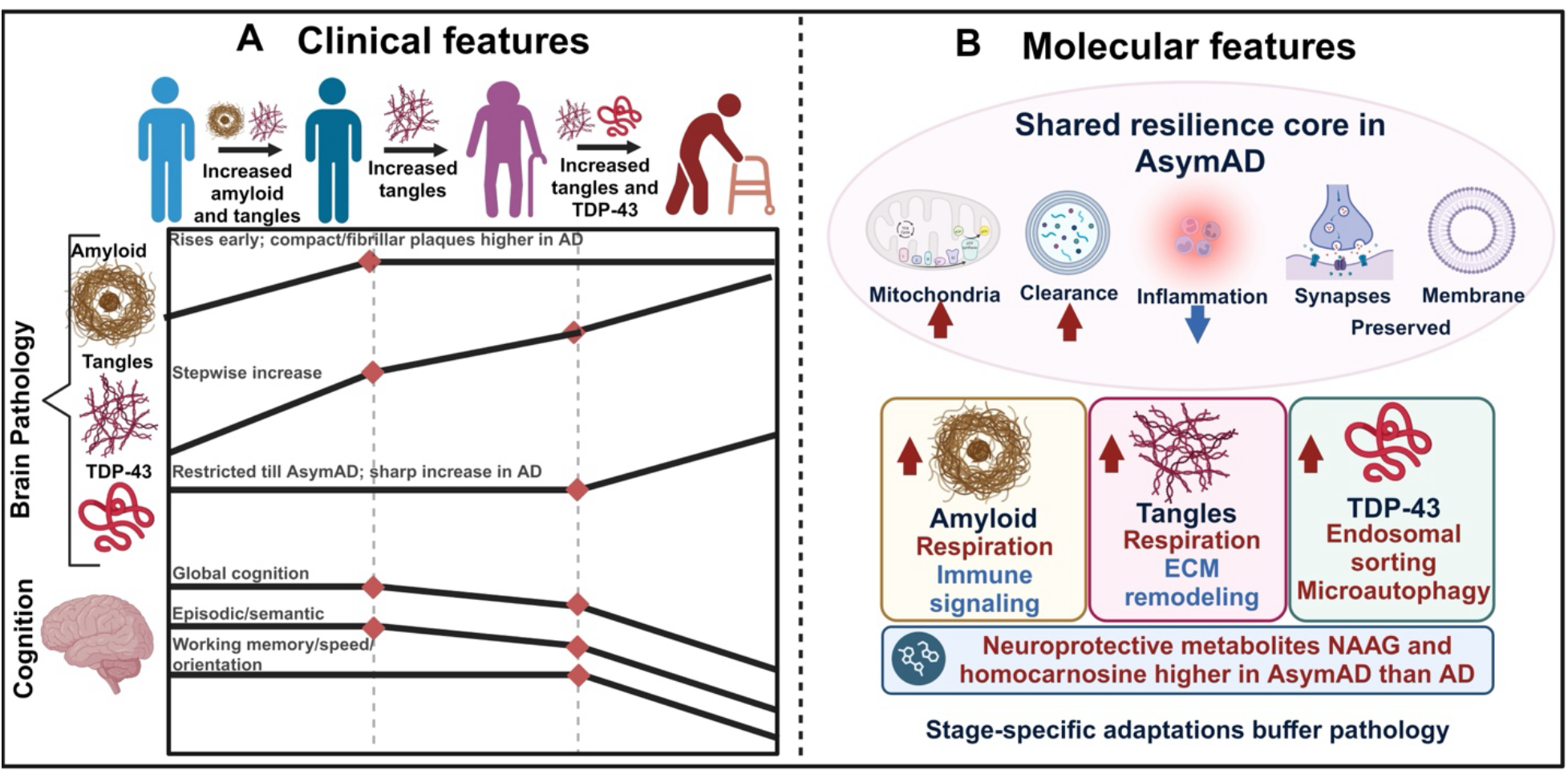
Summary model of resilience in AsymAD. (A) Clinical features: Schematic trajectories of pathology and cognition from controls through resilient and vulnerable AsymAD to AD. Amyloid rises early, tangles increase stepwise, and TDP-43 rises only in AD. Episodic and semantic memory decline in vulnerable AsymAD and AD. (B) Molecular features: AsymAD shows a shared resilience core of higher mitochondrial and endolysosomal clearance proteins, lower inflammation, and preserved synaptic proteins and membrane lipids. Pathology-specific adaptations accompany amyloid, tau, and TDP-43, and NAAG and homocarnosine are higher than in AD. Arrows and colors indicate higher (red) or lower (blue) levels in AsymAD than in AD.

First, AsymAD is not simply AD with less pathology. Earlier studies generally matched AsymAD and AD on CERAD and Braak stage and attributed the remaining clinical difference to protective biology^13,33^. Our analyses suggest that part of the difference lies in the pathology itself. Total amyloid was comparable between AsymAD and AD across SEA-AD regions, yet dense-core and fibrillar plaques were enriched in AD in prefrontal, temporal, hippocampal, and occipital regions, and regional tau was higher in AD. Phosphorylated TDP-43 was concentrated in AD hippocampus and entorhinal cortex. In ROSMAP, TDP-43 was associated with tangle burden but not with amyloid, in agreement with the frequent co-occurrence of LATE-NC with advanced neurofibrillary pathology^2,21,22^. AsymAD individuals therefore appear to tolerate amyloid with less progression to compact plaques, cortical tau, and TDP-43 pathology. This matters for anti-amyloid therapy and prevention trials, because individuals with equivalent amyloid burden may differ in plaque composition and in co-pathology that is currently invisible during life^2^.

More than half of the individuals who were cognitively unimpaired at their final evaluation carried AD-level pathology. Dividing AsymAD by cognitive outcome showed that vulnerability coincided with tangle burden, not with amyloid or TDP-43, and with selective decline in episodic and semantic memory. Tangles begin in the medial temporal lobe and spread to the lateral temporal cortex^34^, regions that support these memory domains. The domain-specific pattern is consistent with earlier longitudinal studies, in which semantic and episodic memory declined years before dementia diagnosis^35,36^ and speeded tasks showed early change^37^. Because our trajectories are aligned to death and groups were defined at autopsy, they describe decline retrospectively and do not test prediction. Whether tracking these cognitive domains together with plasma markers of tau pathology can identify amyloid-positive individuals at risk will require prospective cohorts.

The molecular signature that distinguishes AsymAD from AD centered on preserved mitochondrial translation, one of the most consistent features across all six pathology strata. Mitochondrial dysfunction has long been proposed as an early driver of sporadic AD^38^, and early changes in energy metabolism have been reported in large-scale AD proteomes^39^. Our data indicate that mitoribosomal and translation-factor proteins (TUFM, LRPPRC, SLIRP) remain higher in AsymAD than in AD irrespective of the type and stage of proteinopathy. TDP-43 localizes to mitochondria and impairs the expression of mitochondrially encoded complex I subunits^40^, so restricted TDP-43 pathology in AsymAD may contribute to this preservation, a hypothesis our cross-sectional data cannot test. In the correlation network, mitochondrial translation proteins were coupled positively to homocarnosine and betaine and negatively to 2-aminoadipate, a lysine-degradation intermediate and marker of protein oxidation, and to acisoga, a polyamine oxidation product, which links these metabolites to the bioenergetic program.

The neuronal and lipid components of the signature were also consistent with the literature. NRN1 has been linked to cognitive resilience^18^, somatostatin-expressing interneurons are among the earliest lost in AD cortex^41^, and NAAG is an abundant neuronal peptide that restrains glutamate release^27^. Higher NAAG with lower glutamate is consistent with restrained excitatory signaling, and higher EEA1 suggests preservation of an early endosomal compartment that is enlarged early in sporadic AD^42^. The metabolome added membrane phospholipids: polyunsaturated phospholipids were higher in AsymAD, whereas their breakdown products GPC and glycerophosphoethanolamine were lower. Phospholipase A2, which hydrolyzes these lipids, contributes to cognitive deficits in AD models^43^, and DHA-derived neuroprotective mediators are depleted in AD hippocampus^44^, so lower phospholipid breakdown in AsymAD is consistent with reduced membrane damage. Several of these metabolites, including NAAG, GPC, and myo-inositol, are detectable in vivo by magnetic resonance spectroscopy^45^, which raises the possibility of monitoring this program during life.

Stratification by proteinopathy and stage showed that this shared signature is accompanied by pathology-specific programs. Under frequent amyloid, AsymAD showed lower immune and class I MHC antigen-presentation pathways, in line with glial activation as a correlate of symptomatic disease^46^. Under neocortical tau, it showed lower extracellular matrix and neutrophil degranulation pathways with higher respiration, and under neocortical TDP-43, lower splicing with higher late-endosomal microautophagy and ESCRT-mediated trafficking, which could reflect a compensatory clearance response. These adaptations were identified in strata of modest size, and the amyloid-specific signature grew at the high stage whereas those for tau and TDP-43 did not, so they require validation in larger cohorts. Together, they suggest that resilience combines a common molecular state with responses adapted to the lesion encountered.

A central observation was that the mitochondrial, synaptic, and immune differences between AsymAD and AD were largely confined to AD. Compared with controls, only 10 proteins in resilient AsymAD and 19 in vulnerable AsymAD differed, nearly all higher and related to amyloid deposition or the extracellular matrix (MDK, SMOC1, APP, CTHRC1, NTN1), consistent with the association of these matrisome proteins with amyloid^47^. Resilience may therefore be better described as the maintenance of a control-like molecular state despite amyloid than as an enhanced protective program. Notably, vulnerable AsymAD, which carried more tangles and greater memory decline, did not differ from resilient AsymAD in any protein, and only five proteins separated vulnerable AsymAD from AD, three of them mitochondrial (LRPPRC, TUFM, SLC25A46). This suggests that mitochondrial translation proteins are among the molecular changes closest to the transition from MCI to dementia, although the small number of significant proteins calls for caution. Several factors could explain the absence of proteomic differences between resilient and vulnerable AsymAD. The DFC is involved later than the temporal memory circuits^34^, the groups are modest in size (77 and 65 donors), and bulk tissue may mask cell-type-specific changes.

In summary, cognitive resilience to AD pathology involves a different pathological composition, with less compact amyloid, tau, and TDP-43, and the maintenance of mitochondrial, synaptic, and lipid homeostasis that is lost in AD. Therapeutic strategies may therefore need to combine reduction of the pathologies that distinguish AD from AsymAD, particularly tau and TDP-43, with support of these homeostatic programs. Experimental systems that manipulate mitochondrial translation and phospholipid turnover in the presence of tau and TDP-43 pathology will be needed to test whether these programs causally protect cognition.

### Limitations of the study

This study has several limitations. All molecular data are cross-sectional and postmortem, so the ordering of groups reflects differences between individuals and does not establish temporal or causal sequence within a person. Proteomic and metabolomic profiles were measured in bulk dorsolateral prefrontal cortex, where differences in neuronal and glial content may contribute to the observed changes, and region-resolved pathology was available only in SEA-AD, whose sample size is modest. Metabolites were measured on an untargeted platform and are reported as standardized values, so effect sizes are not comparable with proteomic fold changes. The protein–metabolite correlation network pools AsymAD and AD donors and therefore reflects covariation across these groups rather than regulation within each group. Several pathology strata, particularly neocortical TDP-43 in AsymAD, contained few donors, and the resilient and vulnerable AsymAD groups (77 and 65 donors) may be underpowered to detect small proteomic differences. ROSMAP and SEA-AD participants are predominantly non-Hispanic white and highly educated, which may limit generalizability. Validation in independent and more diverse cohorts, in additional brain regions including the temporal lobe, and in experimental systems will be needed to determine whether these programs causally determine resilience.

## Methods

### Experimental model and study participant details

#### Data acquisition

Data were obtained from participants in the Religious Orders Study (ROS) and Rush Memory and Aging Project (MAP), collectively referred to as ROSMAP. ROS enrolls older Catholic clergy from across the United States, whereas MAP enrolls older adults primarily from the Chicago metropolitan area. Longitudinal cognitive assessments and neuropathological measures were obtained from the Rush Alzheimer’s Disease Center (RADC). Independent regional neuropathological data were obtained from the Seattle Alzheimer’s Disease Brain Cell Atlas (SEA-AD) through the Allen Institute data portal^25^. Proteomic and metabolomic datasets for BA9 (DFC) were obtained from the AD Knowledge Portal hosted on Synapse (syn51196652; syn26007830). Tandem mass tag (TMT) mass spectrometry proteomic data from BA9 were obtained after batch correction and normalization by tunable approach for median polish of ratio (TAMPOR)^16,17^. Values represent log2 relative abundance. Untargeted brain metabolomic data (Metabolon HD4) were processed as described previously^48^. Metabolites with more than 25% missing values were removed (667 of 1,055 retained). Samples were normalized by probabilistic quotient normalization and log2-transformed, outlier samples were removed using the local outlier factor, extreme values were set to missing, and missing values were imputed by k-nearest neighbors. For the analyses reported here, each metabolite was then z-scored, so metabolomic effect sizes are standardized mean differences in SD units. Compounds that Metabolon annotated as lower-confidence identifications are marked with an asterisk.

#### Donor classification

Donors were classified as Control, asymptomatic Alzheimer’s disease (AsymAD), or Alzheimer’s disease (AD) using CERAD neuritic plaque scores^49^, Braak stages^34^, and clinical criteria adapted from Johnson *et al. (2020).* Controls were defined as individuals without dementia at the final clinical evaluation and with CERAD scores of 0-1 and Braak stages of 0-3; for individuals with Braak stage 3, a CERAD score of 0 was required. AsymAD donors were defined as individuals without dementia despite substantial AD neuropathology, with CERAD scores of 1-3 and Braak stages of 3-6. AD donors had dementia at the final clinical evaluation and CERAD scores of 2-3 and Braak stages of 3-6. For ROSMAP, clinical and neuropathological classifications were matched to proteomic data from BA9, yielding 321 donors comprising 79 Controls, 142 AsymAD, and 100 AD cases. The SEA-AD cohort comprised 6 Controls, 27 AsymAD, and 37 AD donors. For SEA-AD, dementia status was defined using the binary dementia annotation provided by the study.

#### Resilient and vulnerable AsymAD

To capture heterogeneity within AsymAD, donors were further classified by their final clinical diagnosis before death. AsymAD donors with no cognitive impairment (NCI) at the final evaluation were classified as resilient AsymAD (n = 77), and those with a final diagnosis of mild cognitive impairment (MCI) were classified as vulnerable AsymAD (n = 65). Together with Controls (n = 79) and AD (n = 100), these groups were ordered as Control → resilient AsymAD → vulnerable AsymAD → AD for trend analyses.

#### Cognitive measures and longitudinal trajectories

Global cognition and five cognitive domains (episodic memory, semantic memory, working memory, perceptual speed, and perceptual orientation) were obtained for ROSMAP annual assessments as standardized composite scores from RUSH. Time was expressed relative to death and restricted to the 20 years preceding death; years before death were calculated as age at assessment minus age at death (−20 to 0). Age at death was mean-centered. For visualization, individual trajectories were plotted with diagnosis-specific LOESS curves (span 0.9) and 95% confidence intervals. Rates of cognitive change were estimated with linear mixed-effects models in lme4 and lmerTest using the BOBYQA optimizer (Bates et al., 2015):

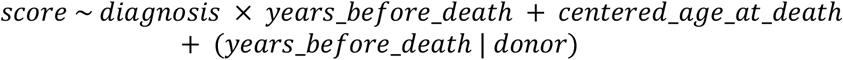

Models included donor-specific random intercepts and slopes, and *P* values used Satterthwaite degrees of freedom. Control served as the reference diagnosis; the Control slope corresponded to the main effect of time, and AsymAD and AD slopes were calculated as the time main effect plus the respective interaction. Diagnosis-specific slopes and Tukey-adjusted pairwise slope comparisons were estimated with emmeans::emtrends. The same framework was applied to global cognition and each domain. Domain-level interaction *P* values in source-data tables were corrected by the Benjamini–Hochberg (BH) procedure. Global cognition was additionally plotted by follow-up year from baseline as group means ± s.e.m., excluding follow-up years with fewer than 10 observations per group.

#### Neuropathological profiling

ROSMAP measures included CERAD score, Braak stage, global AD pathology, regional amyloid burden (amylsqrt_est_8reg), regional tangle burden (tangsqrt_est_8reg), TDP-43 stage (tdp_st4), regional TDP-43 burden (tdp_cs_6reg), and global cognition at the final assessment (cogn_global_lv). Amyloid and tangle burden are square-root-transformed composites across eight regions (angular gyrus, anterior cingulate cortex, calcarine cortex, entorhinal cortex, hippocampus, inferior temporal cortex, midfrontal gyrus, and superior frontal cortex). TDP-43 stage follows the ROSMAP four-level scheme corresponding to LATE-NC stages (Nelson et al., 2019). SEA-AD quantitative neuropathology was used to evaluate 11 regions: angular gyrus (AnG), head of caudate (CaH), dorsolateral prefrontal cortex (DFC), extrastriate occipital cortex (ESOC), frontoinsula (FI), hippocampus (HIP), inferior temporal gyrus (ITG), medial entorhinal cortex (MEC), middle temporal gyrus (MTG), superior temporal gyrus (STG), and primary visual cortex (V1C). Amyloid was quantified as 6E10-positive total, dense-core, fibrillar, and diffuse plaque area; tau as AT8-positive area; and TDP-43 as phosphorylated TDP-43 (pTDP-43) area where available (Gabitto et al., 2024; Travaglini et al., 2026). For group summaries, the mean ± s.e.m. was calculated within each diagnosis and z-scored across diagnoses for each measure to visualize relative burden; annotations show unscaled values. Group differences were assessed by Kruskal–Wallis tests followed by Dunn’s post hoc comparisons with BH correction. AsymAD-AD comparisons within diagnosis and stage strata used two-sided Wilcoxon rank-sum tests with BH correction. Pairwise inter-pathology relationships were estimated by group-wise linear regression.

#### Differential abundance analysis

Differential abundance between AsymAD and AD was assessed for each protein and metabolite using limma (Ritchie et al., 2015) with AD as the reference level:

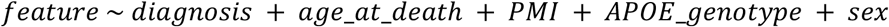

Empirical Bayes moderation was applied, *p* values were BH-corrected, and features with adjusted *p* < 0.05 were considered differentially abundant. Pathway enrichment against the Reactome library was performed with Enrichr^50^.

#### Pathology-stratified analyses

To determine whether AsymAD-AD differences depended on proteinopathy type and stage, donors were stratified into six strata: moderate (CERAD 2) and frequent (CERAD 3) amyloid; limbic (Braak III–IV) and neocortical (Braak V) tau; and early (LATE-NC stages 1–2) and neocortical (stage 3) TDP-43. Within each stratum, AsymAD and AD were compared using the limma model above, and features with nominal *P* < 0.05 were considered stratum-significant. Features unique to each proteinopathy at each stage were identified from Venn intersections. Robust proteins were defined as those significant in the same direction in all six strata (19 higher and 3 lower in AsymAD). Because of smaller metabolomic strata, robust metabolites were defined as those significant in the same direction in at least three strata.

#### Protein-metabolite correlation network

For the 166 donors with matched proteomic and metabolomic profiles (93 AsymAD, 73 AD), we computed Spearman correlations between all differentially abundant proteins (n = 799) and metabolites (n = 53) and corrected the P values across all pairs by the Benjamini–Hochberg method. The network in Figure 3E retains edges with |ρ| ≥ 0.5 and adjusted P < 0.05. To test sensitivity to this cutoff, we recomputed the network at |ρ| ≥ 0.45 and ≥ 0.40. All correlations at |ρ| ≥ 0.40 had adjusted p < 0.001. Lowering the cutoff from 0.5 to 0.4 increased the network from 110 to 809 edges, from 44 to 201 proteins, and from 13 to 34 metabolites (344 edges, 108 proteins, and 23 metabolites at 0.45). GPC and betaine were among the three most connected metabolites, and NAAG among the four most connected, at every cutoff. Homocarnosine rose from 8 protein partners at 0.5 to 92 at 0.4. Correlation coefficients and adjusted P values for all protein–metabolite pairs are in Table S3.

We examined the |ρ| ≥ 0.40 network for subnetworks that correspond to processes highlighted in Figures 3(B-D): mitochondrial translation and membrane phospholipid metabolism. The mitochondrial module contains mitoribosomal subunits (MRPL*, MRPS*, DAP3, PTCD3) and mitochondrial RNA-processing, import, and assembly factors (SLIRP, LRPPRC, ATAD1, ACADSB, the TIMM family, TRMT10C, and the NDUFAF family), with their edges to homocarnosine, betaine, 2-aminoadipate, and acisoga (Figure S4A). The phospholipid module contains proteins correlated with at least one PC or PE carrying a 20:4 or 22:6 acyl chain, with their edges to these phospholipids and to GPC and glycerophosphoethanolamine (Figure S4B). For proteins correlated with both an intact phospholipid and a breakdown product (GPC or glycerophosphoethanolamine), we compared the sign of the mean correlation for the two groups. Figure 3E was drawn in Cytoscape v3.10.4^51^. The networks in Figure S4 were drawn in R (v4.5.2) with igraph (v2.1.2) and ggraph v2.2.2 using a Fruchterman-Reingold layout and a fixed random seed, and two-node components were omitted.

#### Trend analysis across the resilience continuum

Ordered trends across Control, resilient AsymAD, vulnerable AsymAD, and AD were tested for each protein and metabolite with the two-sided Jonckheere–Terpstra test (Jonckheere, 1954), with BH correction across features; adjusted *P* < 0.05 defined increasing or decreasing features. Concordance between the stage-invariant AsymAD signatures and trend direction was summarized as Venn intersections (Figure 5C, 5E). Group means ± s.e.m. are shown for selected features; individual donors are shown in the background.

### Quantification and statistical analysis

All analyses were performed in R (v4.5.2). Statistical tests, sample sizes (n, number of donors), definitions of center and dispersion, and multiple-testing corrections are reported in the figure legends and the sections above. Unless otherwise stated, tests were two-sided, *P* values were corrected by the BH procedure, and adjusted *p* < 0.05 was considered significant. Boxplots show the median and interquartile range, with whiskers extending to 1.5× the interquartile range.

## Acknowledgement

The results published here are in whole or in part based on data obtained from the AD Knowledge Portal (NIH-NIA, U24AG061340), Rush Alzheimer’s Disease Center and Seattle-AD (SEA-AD).

## Code availability

Codes used for analysis are available in the following Github repository: https://github.com/BaloniLab/Proteinopathies_in_Alzheimer-s_Disease. Codes can also be accessed at Mandal, P., & Baloni, P. (2026). Brain proteinopathies associated physiological changes leading to cognitive resilience in Alzheimer’s disease. Zenodo. https://doi.org/10.5281/zenodo.24020520

