## Supplementary figure for "Brain proteinopathies associated physiological changes leading to cognitive resilience in Alzheimer’s disease"

### Supplementary figures

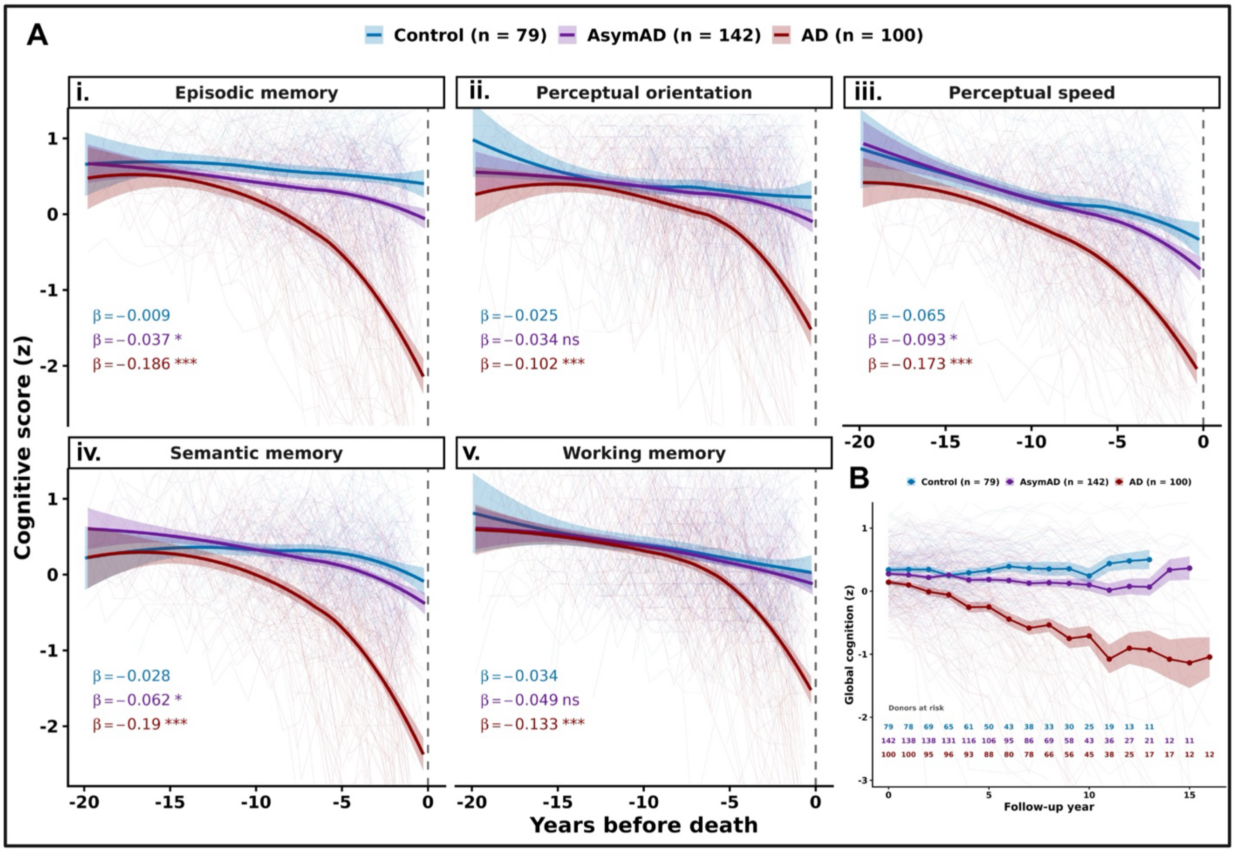


**Figure S1. Domain-specific cognitive trajectories distinguish AsymAD from AD.** (A) Trajectories of individual cognitive domains in ROSMAP as a function of years before death: (i) episodic memory, (ii) perceptual orientation, (iii) perceptual speed, (iv) semantic memory and (v) working memory. Control (blue, n = 79), AsymAD (purple, n = 142), AD (red, n = 100). Thin lines show individual donor trajectories; thick lines show LOESS-smoothed group trajectories (span = 0.9) with shaded 95% confidence intervals. β, rate of cognitive change per year before death, estimated by a linear mixed-effects model with diagnosis × time, age at death (mean-centered), and a random intercept and slope per donor; asterisks indicate the p value (Satterthwaite t-test) for the difference between each group's slope and Control's (reference group, unmarked): $^{*}p < 0.05$, $^{**}p < 0.01$, $^{***}p < 0.001$. (B) Global cognition by follow-up year (years since baseline) for the same groups. Points show group means and shading shows (s.e.m./95% CI); thin lines are individual donors. Donor numbers are mentioned below the plot.

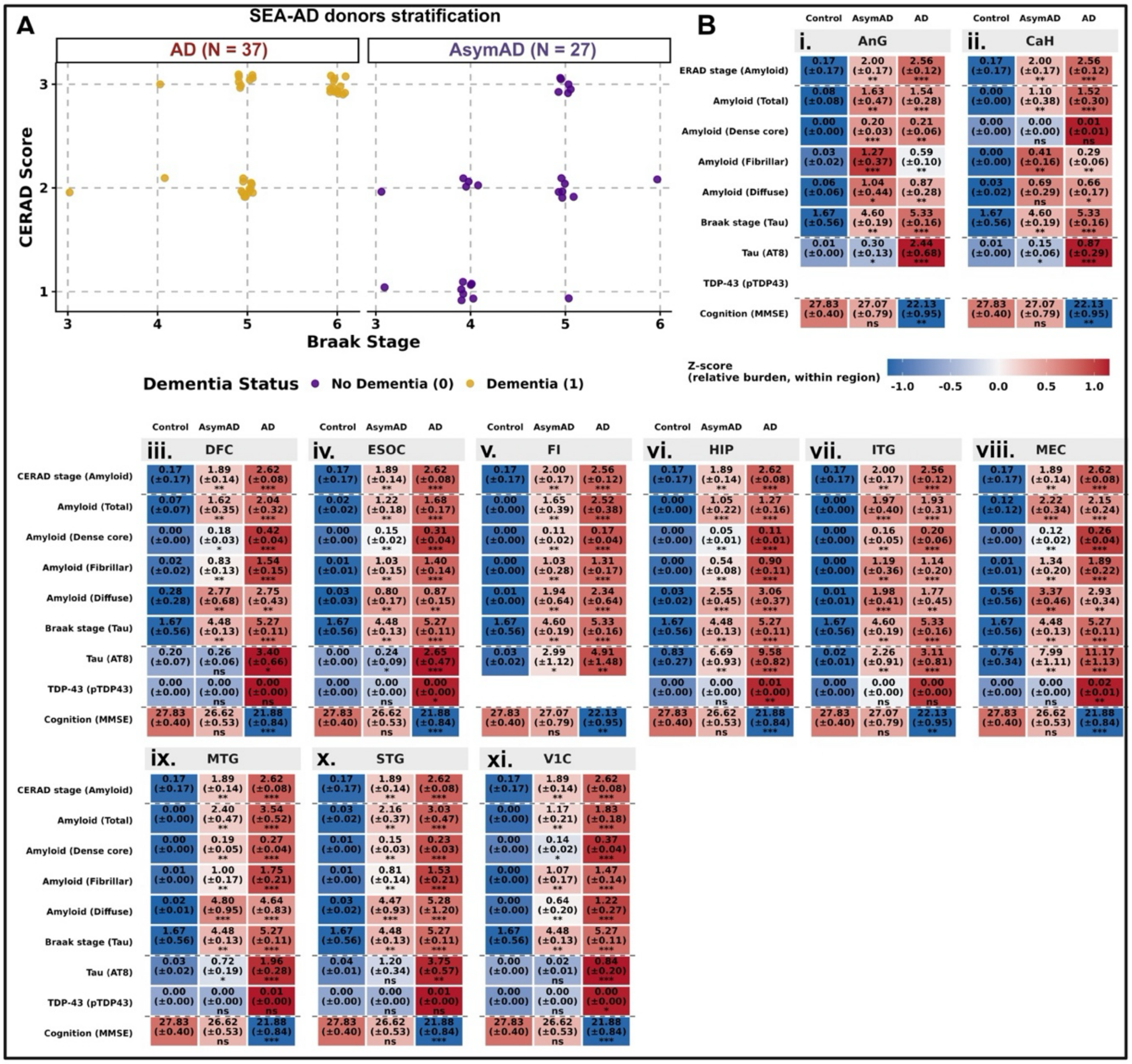


**Figure S2. Stratification and regional neuropathology of SEA-AD donors.** (A) Distribution of SEA-AD donors into AD (n = 37) and AsymAD (n = 27) by Braak stage (x-axis) and CERAD score (y-axis), colored by dementia status (purple, no dementia; yellow, dementia). Points are jittered to reduce overlap. (B) Regional neuropathology and cognition across Control, AsymAD and AD donors in (i) angular gyrus (AnG), (ii) head of the caudate (CaH), (iii) dorsolateral prefrontal cortex (DFC), (iv) extrastriate occipital cortex (ESOC), (v) frontoinsula (FI), (vi) hippocampus (HIP), (vii) inferior temporal gyrus (ITG), (viii) medial entorhinal cortex (MEC), (ix) middle temporal gyrus (MTG), (x) superior temporal gyrus (STG) and (xi) primary visual cortex (V1C). Rows show CERAD stage, quantified amyloid (total, dense-core, fibrillar, diffuse), Braak stage, phospho-tau (AT8), phospho-TDP-43 (pTDP43) and cognition (MMSE). Cell values are mean (± s.e.m.) rounded to 2 decimal places; color shows the within-region z-score of relative burden (scale, upper right). Blank cells indicate measures not available for that region (pTDP43 in AnG, CaH and FI). CERAD, Braak and MMSE are donor-level and repeated across regions. Asterisks denote significance versus Control by Dunn's test, Benjamini-Hochberg correction: ns, not significant; *$p < 0.05$, **$p < 0.01$, ***$p < 0.001$.

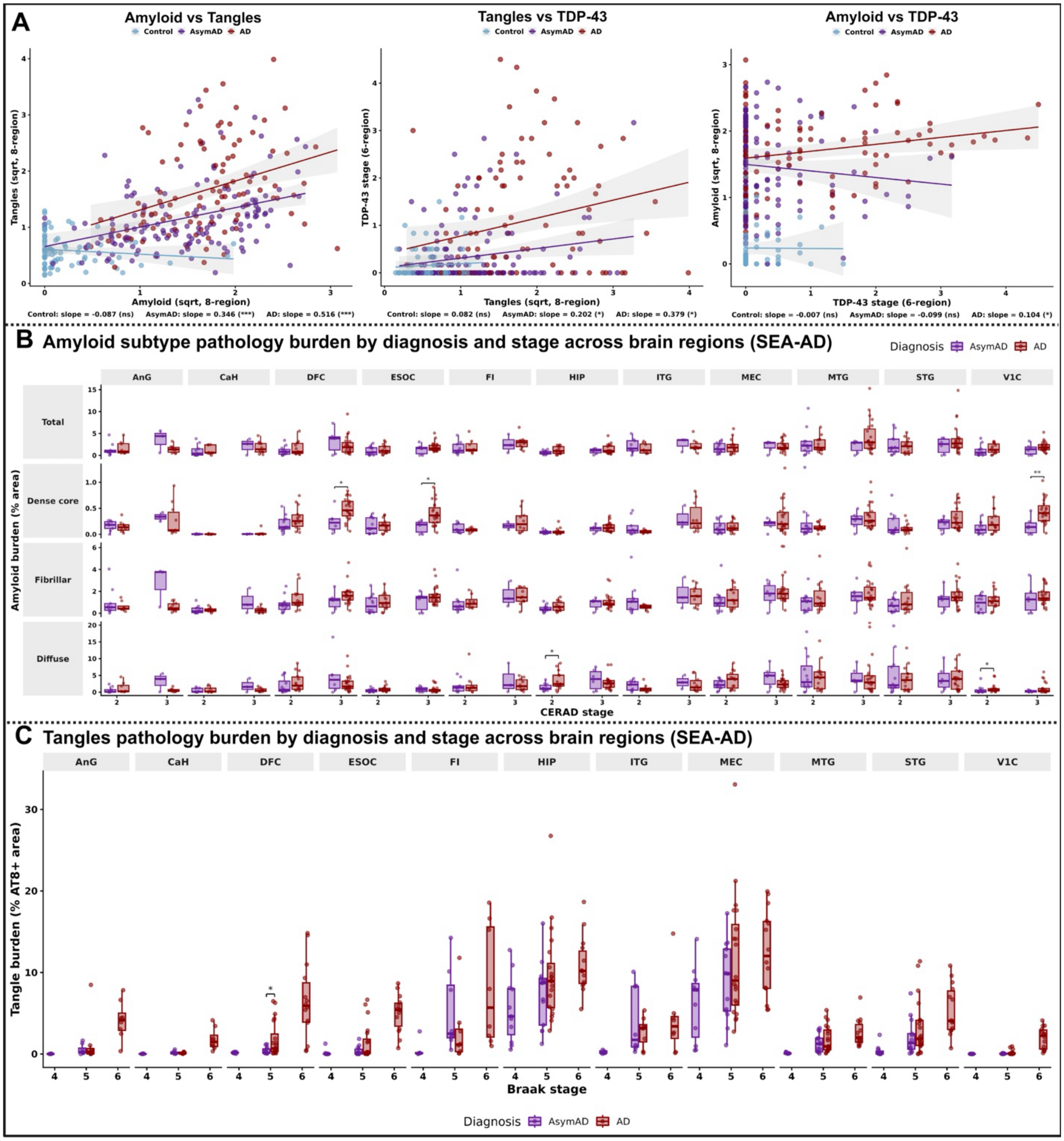


**Figure S3. Inter-pathology relationships and stage-matched regional comparisons of AsymAD and AD.** (A) Pairwise relationships among amyloid, tangle and TDP-43 pathology in ROSMAP: amyloid versus tangles (8-region, square-root transformed), tangles versus TDP-43 stage (8-region), and TDP-43 stage versus amyloid (8-region, square-root-transformed). Each point is one donor (Control, blue; AsymAD, purple; AD, red). Lines show [group-wise linear regression fits] with shaded 95% confidence intervals. Slopes and their significance are given below each panel: ns, not significant; $*P < 0.05$, $***P < 0.001$. (B) Amyloid subtype burden (total, dense-core, fibrillar and diffuse; percent 6E10-positive area) in SEA-AD AsymAD (purple) and AD (red) donors, stratified by CERAD stage (2 or 3) within each of 11 brain regions (AnG, CaH, DFC,

ESOC, FI, HIP, ITG, MEC, MTG, STG, V1C). Boxes show median and IQR; points are individual donors. (C) AT8-positive tangle burden (percent area) in SEA-AD AsymAD and AD donors stratified by Braak stage (4–6) within each region. Boxes show median and IQR; points are individual donors. Brackets indicate AsymAD versus AD within a stage by two-sided Wilcoxon rank-sum test, Benjamini–Hochberg-adjusted: *$p < 0.05$, **$p < 0.01$.*

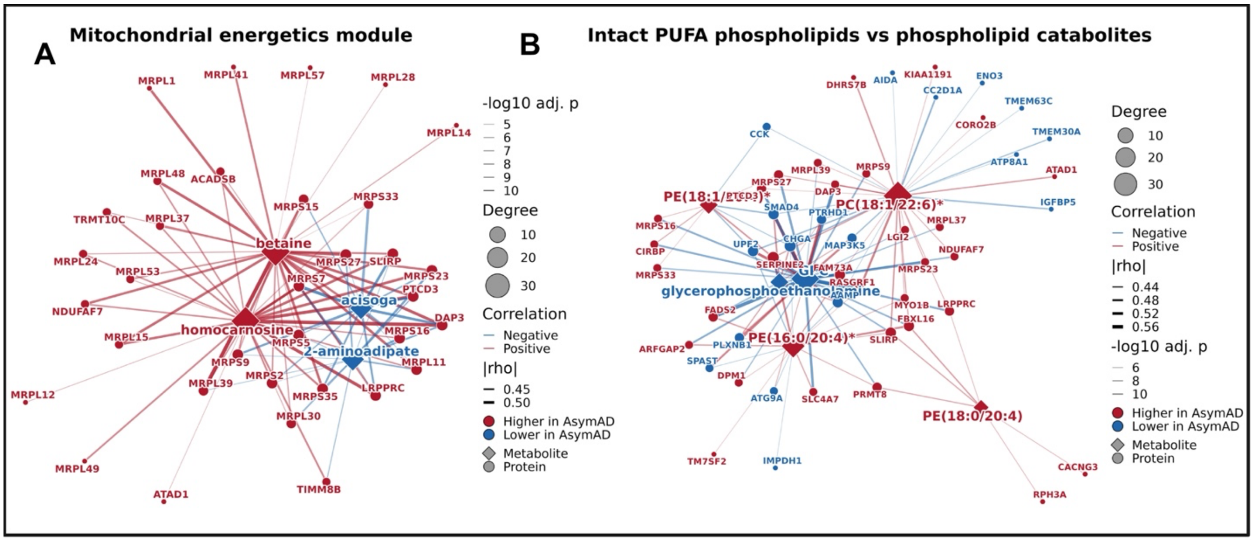


**Figure S4. Protein–metabolite correlation network resolves mitochondrial energetics and phospholipid modules distinguishing AsymAD from AD**. (A) Mitochondrial energetics module at $|\rho| \geq 0.40$. The panel shows correlations of mitochondrial proteins (mitoribosomal subunits and mitochondrial RNA-processing and assembly factors) with homocarnosine, betaine, 2-aminoadipate, and acisoga. (B) Phospholipid module at $|\rho| \geq 0.40$. Proteins correlated with at least one PUFA-containing phosphatidylcholine (PC) or phosphatidylethanolamine (PE) with 20:4 or 22:6 acyl chains are shown. Their edges to these intact phospholipids and to the phospholipid catabolites glycerophosphorylcholine (GPC) and glycerophosphoethanolamine are included. Acyl chain composition is shown as sn-1/sn-2. Isolated two-node components were omitted for clarity. In (A) and (B), diamonds represent metabolites and circles represent proteins. Node color indicates direction of change (red, higher in AsymAD; blue, lower in AsymAD), and node size is proportional to degree. Edge color indicates positive (red) or negative (blue) correlation, edge width is proportional to $|\rho|$, and edge opacity reflects −log10 adjusted p value. Correlations were computed across 166 donors with matched BA9 proteomic and metabolomic profiles (93 AsymAD, 73 AD). They were restricted to the 799 proteins and 53 metabolites differentially abundant between AsymAD and AD, and p values were adjusted by the Benjamini–Hochberg method. Metabolites marked with an asterisk denote compounds identified with lower confidence.

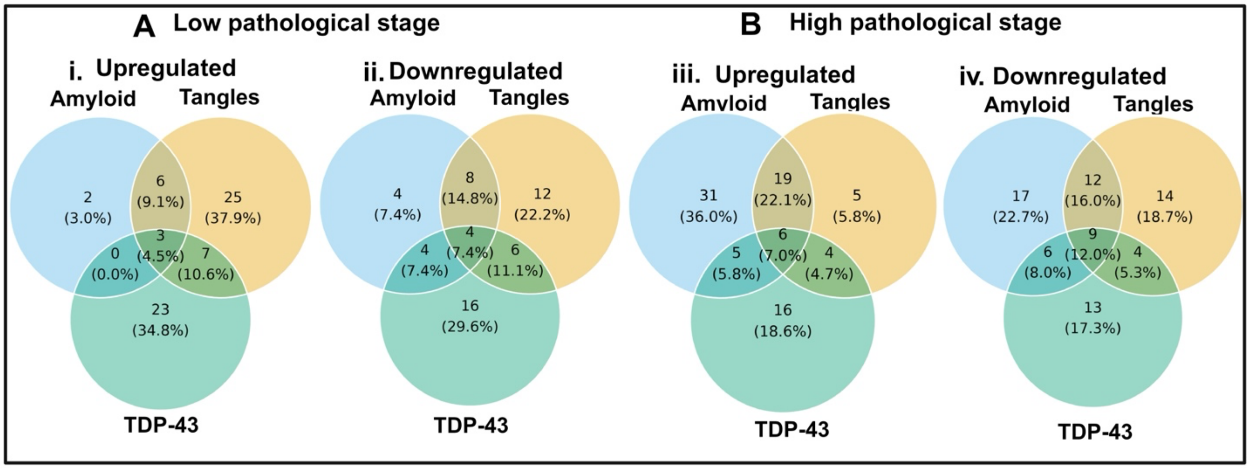


**Figure S5. Stage-stratified overlap of differentially abundant metabolites**. (A and B) Venn diagrams of metabolites differentially abundant between AsymAD and AD in the DFC (BA9) within amyloid, tau, and TDP-43 strata at the low (A) and high (B) pathological stage. Metabolites higher (i, iii) and lower (ii, iv) in AsymAD are shown separately. Values are the number of metabolites in each region, with the percentage of all metabolites in the diagram in parentheses. Low-stage strata were moderate amyloid (CERAD 2; n = 70 AsymAD, 76 AD), limbic tau (Braak III–IV; n = 111, 77), and early TDP-43 (LATE-NC stages 1–2; n = 52, 51). High-stage strata were frequent amyloid (CERAD 3; n = 52, 98), neocortical tau (Braak V; n = 35, 94), and neocortical TDP-43 (LATE-NC stage 3; n = 20, 71).

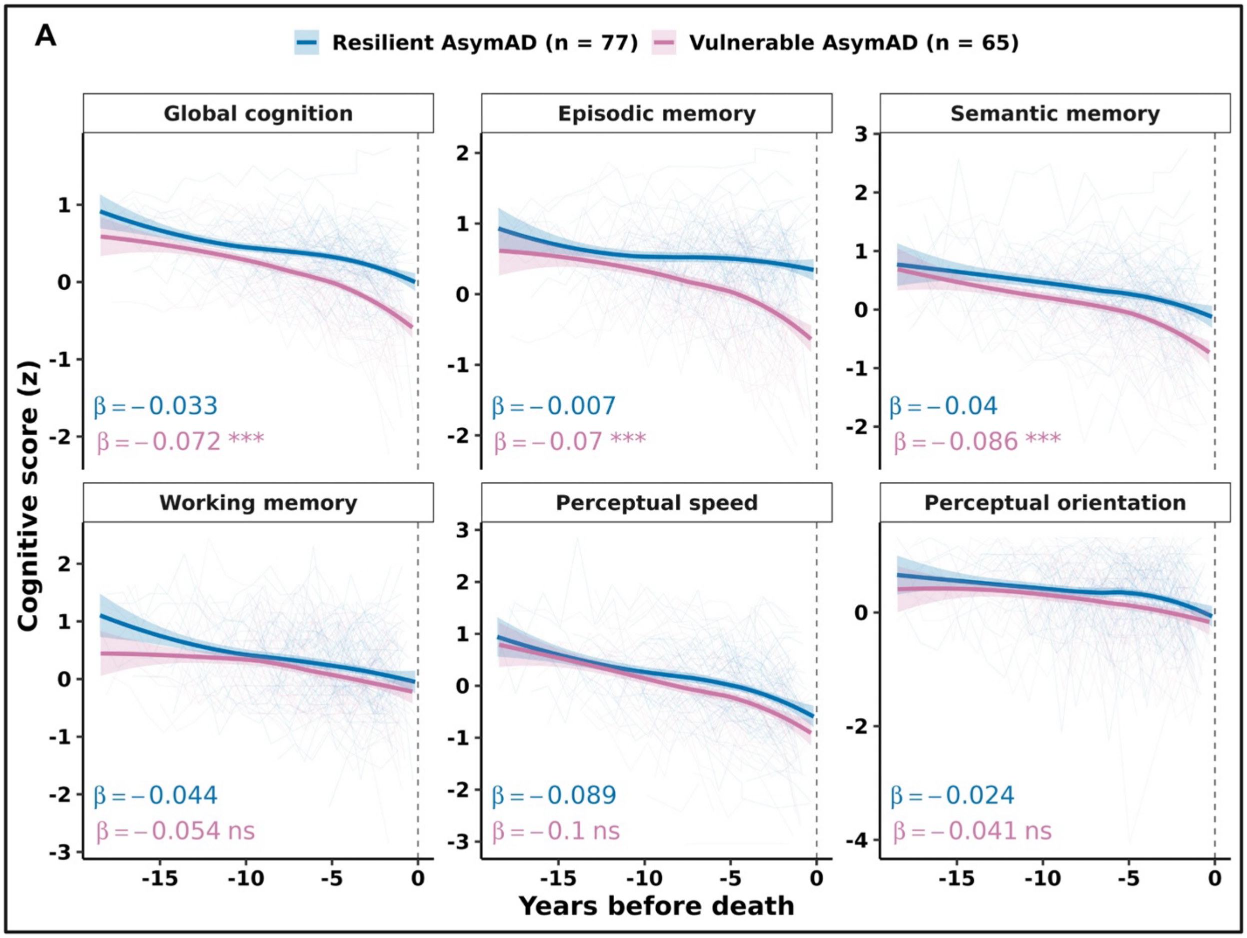


**Figure S6. Domain-specific cognitive trajectories in resilient and vulnerable AsymAD.** (A) Global cognition and five cognitive domains in resilient (blue, n = 77) and vulnerable (pink, n = 65) AsymAD as a function of years before death. Thin lines show individual donors, thick lines the smoothed group trajectory with 95% confidence intervals, and the dashed line death. β is the rate of change per year (SD/year) from a linear mixed-effects model with diagnosis × time, mean-centered age at death, and a random intercept and slope per donor. Asterisks denote a significant difference in slope from resilient AsymAD (interaction p): ***p < 0.001; ns, not significant.

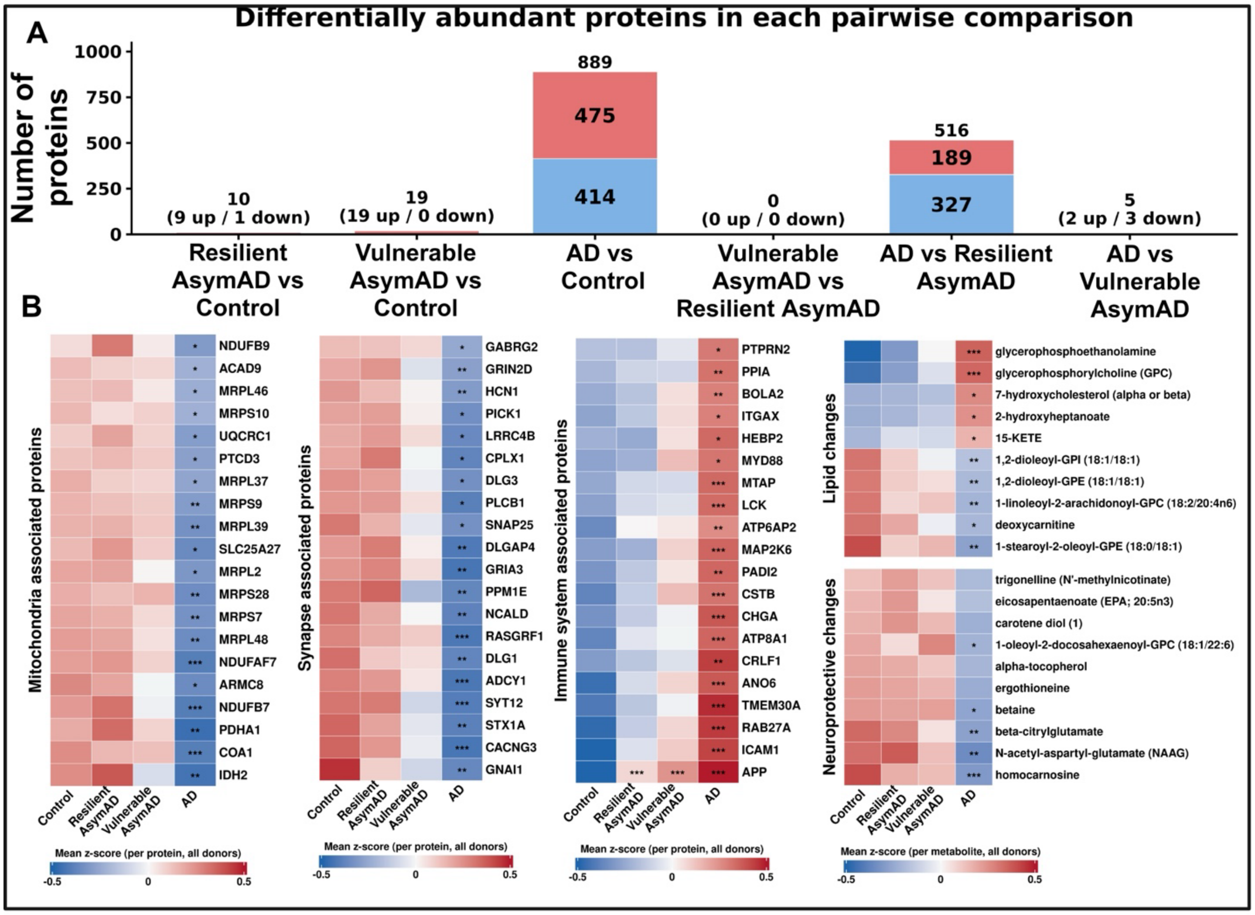


**Figure S7. Pairwise proteomic differences across the AD continuum**. (A) Proteins differentially abundant in each pairwise comparison among controls, resilient AsymAD, vulnerable AsymAD, and AD (limma on all 8,339 quantified proteins, adjusted for age at death, sex, postmortem interval, and *APOE*genotype; adjusted $p < 0.05$). Red indicates higher and blue lower abundance in the first-named group. Statistics for all proteins are in Table S5. (B) Representative features with significant Jonckheere–Terpstra trend. Twenty proteins per module were chosen from proteins annotated to mitochondrial, synaptic, or immune Reactome pathways, requiring AD versus control adjusted $p < 0.05$, a monotonic group trend (Spearman $\rho \geq 0.8$), and ranking by effect size weighted by pathway specificity. For metabolites, the 10 most significant trending lipid metabolites and the 10 most significant trending metabolites from a curated neuroprotective list are shown. Color shows the mean z-score per feature; asterisks show each group versus controls (limma, Benjamini-Hochberg; *adjusted $p < 0.05$, **adjusted $p < 0.01$, ***adjusted $p < 0.001$).
